# Polymorphism can extensively reshape the genome-wide crossover landscape in *Arabidopsis thaliana*

**DOI:** 10.1101/2025.09.19.677288

**Authors:** Benoît Madec, Maëla Sémery, Qichao Lian, Mohamad Yassine, Loïse Léonard-Moniot, Éric Espagne, Sophie Piquerez, Chloé Girard

## Abstract

Meiotic recombination, via the formation of crossovers (COs), re-assorts parental alleles and dictate which traits are inherited together, or not. The recombination rate is not homogeneous across the genome, and uncovering the causes of these fluctuations can inform our understanding of DNA recombination and genome evolution. In this study we show that presence of sequence divergence (polymorphism) between homologous chromosomes increases the local recombination rate in *Arabidopsis thaliana*. This effect is observed genome-wide, turning cold regions of the genome hot. This relocation of COs towards polymorphic regions occurs at the expense of non-polymorphic ones, which become colder. This global overhaul of the CO landscape is dependent on the mismatch repair (MMR) machinery. This suggests that MMR has a pro-CO role, driving COs to regions of the genome where it could detect sequence differences between homologous chromosomes. Our study demonstrates that polymorphism can be a major recombination driver in *A. thaliana*.

## Introduction

Meiotic crossovers (COs) rearrange parental alleles into new combinations on which selection can act. The CO landscape, shaped by the number and the distribution of CO events, dictates which traits will be re-assorted, and which will remain linked. Only few COs are formed per chromosome, despite a 2-200 fold excess of precursors, the DNA double-strand breaks (DSBs). The CO landscape therefore results not only from the underlying DSB distribution, but also from the CO designation process.

Variation in recombination rate (RR) is observed along the genome in all species. Conserved cold regions, with lower RR, include centromeric regions, which are devoid of COs by the combined effect of decreased DSB number and CO suppression (Castellani et al., 2024; Fernandes et al., 2019). In some species including *Arabidopsis thaliana*, humans and the female stickleback, High-Recombining Zones (HRZ) flank this centromeric non/low-recombining zone (NRZ/LRZ), accumulating many COs (Capilla-Pérez et al., 2021; Drouaud et al., 2007; Durand et al., 2022; Fernandes et al., 2024; Giraut et al., 2011; Jing et al., 2024; Kong et al., 2002; Lian, Solier, et al., 2022; Venu et al., 2024).

In *A. thaliana*, this specific pattern of HRZ surrounding centromeric NRZ/LRZ is observed in both sexes as well as in all F1 hybrids tested to date (Capilla-Pérez et al., 2021; Salomé et al., 2012). It also matches the historical CO landscape (Lian, Solier, et al., 2022), showing that it has been a feature of this species throughout its evolution. It also closely follows the distribution of sequence diversity along chromosome length (Alonso-Blanco et al., 2016; Lian, Solier, et al., 2022), indicating a positive association between polymorphism density and CO frequency. This positive association is found in many species (Andersen et al., 2012; Nordborg et al., 1996; J. Wang et al., 2016) and would mainly reflect the fact that recombination shapes the maintenance (in high recombining regions) and the removal (in low recombining regions) of sequence diversity (Charlesworth and Campos, 2014). Supporting this interpretation, in pure lines of *A. thaliana* with virtually no polymorphism between homologous chromosome, COs still accumulate in peri-centromeric HRZ (Lian, Solier, et al., 2022). This indicates that the accumulation of sequence divergence in the peri-centromeric regions is a consequence, rather than a cause, of the higher recombination rate in these regions. This also suggests that other pro-CO forces are driving recombination to peri-centromeric HRZ.

This would argue against a strong impact of polymorphism on CO rate. However, in *A. thaliana*, presence of a polymorphic region in an otherwise non-polymorphic chromosome pair was shown to boost recombination in the polymorphic interval (Blackwell et al., 2020; Ziolkowski et al., 2015). This suggested that presence of polymorphism could be a pro-CO driving force, when competing with non-polymorphic regions. In this study, we systematically characterize the effect of polymorphism on recombination rate along the whole genome of *A. thaliana*.

To do so, we devised a method to specifically test whether COs form preferentially in polymorphic or non-polymorphic intervals, independently of other chromosomal features. This method relies on the use of Recombinant Inbred Lines (RILs), allowing to test this effect at the genome-wide scale (Supplementary Fig. S1). Using this experimental set up, we show that along each chromosome, polymorphic intervals recombine more than non-polymorphic ones, irrespective of their chromosomal position. This pro-CO driving force is strong enough to relocate COs from hot to cold regions. This demonstrates that presence of polymorphism not only favors CO formation genome-wide, but also that it out-competes other pro-CO driving forces that have shaped the CO landscape throughout *A. thaliana* evolutionary history.

The observed reshaping of the CO landscape is not only due to the presence of polymorphism but also to its interaction with CO interference, by which COs tend to be positioned further apart than expected by chance (Girard et al., 2023). Because CO positions along one chromosome are inter-dependent, boosting of recombination in one polymorphic region occurs at the expense of non-polymorphic ones, which become colder all along the genome.

This complete overhaul of the CO landscape is dependent on the mismatch repair (MMR) machinery: in absence of MSH2, COs are favored in homozygous, non-polymorphic, regions rather than heterozygous ones. This points to a pro-CO role of MMR in polymorphic regions, as opposed to the situation in yeast where MMR has an anti-CO role (Cooper et al., 2018; Martini et al., 2011).

Altogether, we propose a model by which the presence of polymorphism, detected by the mismatch repair machinery, together with CO interference can explain many aspects of the CO landscape in *Arabidopsis thaliana*.

## Methods

### Plant material and growth conditions

Recombinant inbred lines from the family 7RV (Col-0 x Ct-1) were provided by the Arabidopsis thaliana stock center of Versailles (http://publiclines.versailles.inra.fr/). These lines were generated from a cross between the 162AV Ct-1 parent and the 186AV Col-0 parent. From this family, two lines were used, 7RV498 and 7RV168 (Supplemental Fig. S2). EMS mutagenesis lines were provided by the team of Raphaël Mercier (Lian, Solier, et al., 2022). Plants were grown at 20°C under long day conditions (16h day, 8h night), in greenhouses or growth chambers from the I2BC greenhouse facility.

### Marker sets elaboration

Frozen leaf material was sent to the Max Planck-Genome-centre to perform library preparations and DNA sequencing (https://mpgc.mpipz.mpg.de/home/). Plants were sequenced at a coverage of 5X (2 × 150bp ; 2.5 million reads). Illumina reads were trimmed using the paired-end option of trimmomatic (v0.39, Bolger et al., 2014). Following trimming, reads were mapped onto the TAIR10 Arabidopsis thaliana reference genome using BWA (v0.7.3, H. Li and Durbin, 2009) and duplicates were removed using sambamba (v1.0, Tarasov et al., 2015). Two different methods were used to establish the marker sets used for downstream CO calling. For variants between Col-0 and Ct-1, variant calling from the CO mapping population was performed with HaplotypeCaller from the GATK suite (v4.2.2.0, Poplin et al., 2017). For EMS markers, we used the method described in Lian, Solier, et al., 2022 and kept only unique, heterozygous SNPs from the different EMS treated lines, from the same accession, using 60X sequencing of the parents unsing the inGAP-family v1.0.0 tool, (Lian, Chen, et al., 2022). For both sets, only variants following a correct Mendelian segregation were kept in the final marker list.

### Genotyping-by-sequencing / CO analysis

These marker sets were used to genotype (inGAP-family v1.0.0) all offspring. Then, two methods were used to detect crossovers, one for heterozygous regions, using Col-0/Ct-1 markers, and one for homozygous regions, using the EMS markers. Both approaches used a sliding-window based method as described in Lian, Solier, et al., 2022. For heterozygous regions, windows of 100Kb, with a step of 50Kb were used, for approximately 400 markers and 1-2,000 reads per window. For homozygous regions, windows were defined by the number of markers, and not physical distance. We used windows of 6 markers, with a step of three markers, irrespective of the genomic distance. In both cases, a CO was validated if the genotype was supported by 5 adjacent windows on each side of the transition, two for chromosome ends. All COs were then checked individually manually.

### Secondary analyses

For heatmaps presented in Figure 2, the number of crossover events was retrieved within every interval and reported to the size of the population. Normalization was then done for each interval using: heterozygous intervals, homozygous intervals and F1 hybrid controls.

The CO events in peri-centromeric regions presented in Fig.5 were retrieved from the 8 long arms peri-centromeric regions, as defined in Underwood et al., 2018 (Supplementary Fig.S3A).

For the analysis of CO distribution at borders presented in Fig.3A and B, all events were retrieved around 1 and 2Mb of the borders. In order to use the same resolution for both homozygous and heterozygous sides, the CO positions were determined using only Col-0 EMS markers, on both sides of the borders. 13 2Mb frontiers and 19 1Mb frontiers were selected across the two RILs (Supplementary Fig.S3B). For a frontier to be selected, the distance between the frontier and the next feature (other frontier, centromeric repeats, end of chromosome) had to be twice as far (more than 2Mb away for 1Mb frontiers, more than 4Mb away for 2Mb frontiers).

For the analysis of patterns along heterozygous intervals presented in Fig.3C, seven distributions of intervals located at chromosome ends were aggregated (Supplementary Fig.S3C). The CO positions were determined using the Col-0 / Ct-1 divergence markers. In order to aggregate the distributions within these intervals, the 114 - 160 windows of 50Kb constituting the 7 intervals were interpolated to 200 windows, using approxfun in R (https://www.rdocumentation.org/packages/stats/versions/3.6.2/topics/approxfun), and then pooled.

For chromosome ends presented at Fig.2C, the number of crossovers formed in the same intervals as Supplementary Fig.S3C plus the right chr-end interval of chromosome 1 of the RIL 7RV498 were used.

Coefficient of coincidence (CoC) curves were obtained as in Girard et al., 2021.

### Statistics

Statistical analyses were done using the PRISM software and R basic packages. The Kruskal-Wallis test was done to compare the number of COs per individual between controls and RIL backcrossed data as well as compare the mean number of COs in heterozygous and homozygous regions, COs at pericentromeres and at chromosome ends. For comparison of WT and *msh2* bRILs, the Mann-Whitney test was done. For double COs and triple COs, a chi-square test was done to compare the number of events associated with the size of the population.

### CRISPR-Cas9 mutagenesis of MSH2 in EMS lines and RILs

Full knock-out and partial knock-out mutants for MSH2 were generated. RNA guides were designed using CRISPOR (https://crispor.gi.ucsc.edu/, Concordet and Haeussler, 2018) and CHOPCHOP (https://chopchop.cbu.uib.no/, Labun et al., 2019). Two guides were selected in 5’ of the TSS and two guides in 3’ of the TTS (Supplementary Table S7). Using goldenbraid as described by Sarrion-Perdigones et al., 2013, gRNA were inserted in plasmids with the Cas9 and a Dsred report gene expressed in seeds (Supplementary Table S8). Plant transformation was done by floral dipping in a *Agrobacterium tumefaciens* solution. Transformed seeds were screened using the Dsred reporter gene. One month after sowing, heat cycles were performed (2 days at 30°C degrees during day and 37°C during night, 3 days at 22°C ; 3 cycles) in order to increase the efficacy of the Cas9 (originally from *Staphilococcus aureus*). At this first T1 generation, plants were screened for full knock-out mutations using PCR primers flanking the position of guide RNAs (Supplementary Table S9). Seeds of T1 mosaic mutants were screened for the loss of the Dsred reporter gene (and the Cas9) and a second genotype screen of T2 mutants was done, followed by Sanger sequencing to confirm the mutation. Two successive backcrosses were done to limit the number of Cas9 offtarget mutations.

## Results

### A method to study the impact of polymorphism on crossover distribution genome-wide

To study the global effect of polymorphism on the genome-wide recombination rate, we took advantage of the genomic structure of Recombinant Inbred Lines (RILs). The genome of a RIL is a homozygous patchwork between two parental genomes. Our strategy (Supplementary Fig. S1) was to backcross a chosen RIL to both its parents to generate two F1 lines, henceforth referred to as bRILs (backcrossed RILs), mirror of each other, where every region of the genome is either present as polymorphic (heterozygous) and non-polymorphic (homozygous). By comparing the two reciprocal bRILs, we can assess whether COs form more frequently in polymorphic or non-polymorphic regions, independently of the genomic location.

Because we want to test the direct *cis* effect of sequence divergence on recombination, without any perturbation from any global *trans* effect (recombination rate modifiers), we chose RILs from the Columbia-0/Catania-1 family (Col-0/Ct-1, 0.4% divergence, ∼1 SNP/250bp), as it was previously shown that no *trans* modifier of the global recombination rate was present between these two ecotypes (Ziolkowski et al., 2015). Within the Col-0/Ct-1 RIL family available from VASC (Ricou et al., 2025), we chose two lines displaying at least one transition between the Ct-1 and Col-0 haplotypes per chromosome, based on genotyping data (Simon et al., 2008, Supplementary Fig. S2). We backcrossed these two RILs to both Col-0 and Ct-1 lines treated with EMS (from Lian, Solier, et al., 2022), to use the few EMS mutations (∼5 SNPs per Mb) as markers to detect CO events within homozygous regions. No meiosis defects could be observed in the bRIL plants (Supplementary Fig. S4), suggesting that recombination proceeded normally. Offspring plants from the bRIL were sequenced to analyze recombination in both male and female meiosis independently (Supplementary Fig. S1).

The position of COs within the different polymorphic and non-polymorphic blocks was detected by using two sets of markers: (i) the natural divergence between Col-0 and Ct-1 in polymorphic regions (411,187 SNPs), and (ii) the EMS markers present in the Col-0 (∼700 SNPs) and Ct-1 (∼400 SNPs) lines in non-polymorphic regions (Supplementary Table S1). In both cases, the detection of CO positions was performed as in Lian, Solier, et al., 2022 (see Methods). To ensure that COs could be efficiently detected with both methods, the CO landscape of F1 hybrids between a Col-0 EMS line and a Ct-1 EMS line was analyzed (Supplementary Fig. S1). In this control cross (comprising 139 female and 145 male meioses), the exact same COs in the same individuals were detected using both sets of markers (Table S3 and S4). The 411,187 Col-0/Ct-1 markers allow for a very precise detection of events, with a mean resolution 5,295 base pairs (Supplementary Table S2). When using the ∼700 Col-0 EMS SNPs, the CO landscape was indistinguishable from the first (Supplementary Fig. S5), indicating that ∼700 SNPs are enough for accurate detection of CO events, albeit with lower resolution (Supplementary Table S2). For the Ct-1 EMS markers, the landscape differs in some regions lacking SNPs, but the global number of COs detected was roughly unchanged (Supplementary Tables S3 and S4). Where the two distributions disagree (*e.g.*, peri-centromeric region on chromosome 4), these regions were not taken into account in their homozygous state (Supplementary Fig. S3). Throughout our data comprising 761 male meioses and 575 female meioses, we detected 5,946 CO events, with a mean resolution of 2.191 Mb (Supplementary Table S2).

In conclusion, we have designed a robust method for the detection of COs in both homozygous and heterozygous regions from sequencing data. Our crossing plan allows for the study of both male and female products of meiosis faithfully.

### In *A. thaliana*, heterozygous regions recombine more than homozygous ones genome-wide

Thanks to our cross scheme, we can probe the number of COs in each individual region, under both their polymorphic and non polymorphic status. For this analysis, both female and male were studied separately, then aggregated for a global description of the effect, and compared to the control F1 hybrid.

**W**e observed the same number of CO events in homozygous regions compared to the F1 hybrid (Fig. 1A, mean= 0.032 CO/Mb per gamete per interval for both F1 hybrid and bRIL homozygous). We observed however a significant increase in CO number in heterozygous regions at the genome-wide scale (Fig. 1A, mean = 0.043 CO/Mb for bRIL heterozygous). This shows that presence of polymorphism can promote recombination in *A. thaliana* globally across the genome. These differences can also be observed in male and female populations (Supplementary Fig. S6C).

Because we performed both reciprocal crosses (to Col-0 and Ct-1), we can compare the crossover rate across the different regions in both situations, polymorphic or non-polymorphic (Fig. 1B and C). Along chromosomes, we observed that the recombination rate profile (cM/Mb) is either similar and most often elevated when the region is heterozygous (purple) compared to when homozygous (yellow). This is true across the whole genome in both females and males (Supplementary Fig. S7 and S8). Moreover, heterozygous regions of the bRILs tend to recombine more than the F1 controls as well (purple v. grey, Supplemental Fig. S7 and S4), although this is not always to case (see below).

This general observation from the smoothed CO landscape is corroborated when looking at the total number of COs per region in the bRILs (Fig. 1D): along each chromosome, heterozygous regions (purple) always show higher recombination frequency (darker color palette) than homozygous ones, independently of their genomic position. This is true in both males and females, and in both RILs tested (Supplementary Fig. S7 and S8). This observation does not hold for the small arm of chromosome 4 however, where a 1.17Mb inversion is present between the Col-0 and Ct-1 genome (Schmidt et al., 2020), confirming that large genome rearrangements prevent CO formation.

Our strategy allowed to show that, in the bRILs, presence of polymorphism is always associated with higher recombination rate, in both male and female meiosis, independently of the genomic location.

### Presence of polymorphism can turn cold regions of the genome hot

In many species, the number of COs per cell is quite stable, with only 10% to 20% variation (S. Wang et al., 2019), and so is the distribution along chromosomes. In *A. thaliana* female meiosis, about half of COs occur in the peri-centromeric regions in the middle of the chromosomes, while sub-telomeric regions are almost devoid of events, in accordance with previous studies (Capilla-Pérez et al., 2021; Giraut et al., 2011).

However, on bRILs chromosomes where large haplotype blocks are present at chromosome ends (Fig. 2A), 4-times more COs were formed when the block was heterozygous in females compared to F1 hybrids (Fig. 2C). However, when a smaller heterozygous region (less than 3Mb) is present at chromosome end, the redistribution was not as strong (Supplementary Fig. S7, Chromosome 3 RIL 7RV168), suggesting that the effect of redistribution could be dependent on the size of the polymorphic region. From these results, we show that the presence of polymorphism in historically low-recombinogenic regions can promote significant CO formation, turning cold regions of the genome hot.

In many species, the CO rate is markedly different in male and female meiosis. This so-called heterochiasmy manifests in *A. thaliana* by an almost two-fold difference in CO number per chromosome, with 1 CO in females and 2 COs in males (Supplementary Fig. S6). The underlying assumption is that in males, chromosome axis length can accommodate for double COs to be formed. However in females, because on one hand axis length is shorter (Drouaud et al., 2007) and interference is stronger (Capilla-Pérez et al., 2021), and on the other hand most COs occur in the middle of the chromosome, this means that no other crossover can be formed on the same chromosome pair, hence preventing double COs to occur. We therefore wondered whether turning sub-telomeric regions hot in females could allow for double COs to be formed.

**Fig. 1.**
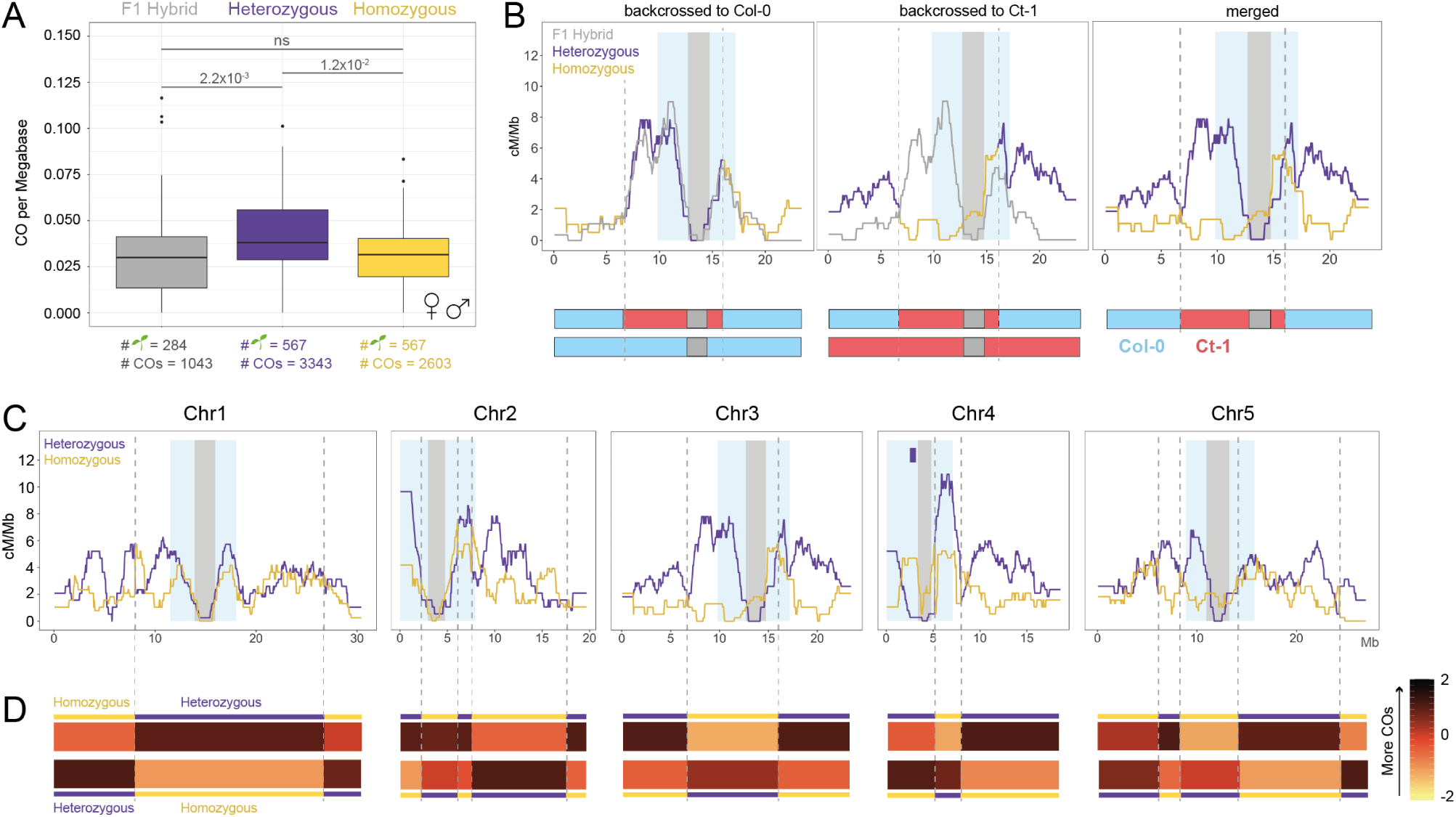
Crossovers preferentially forms in heterozygous rather than homozygous regions. **A)** The normalized number of crossovers per Mb within 35 intervals per plant in aggregated female and male meiosis. Kruskal-Wallis test was used to calculate statistical significance. **B)** The distribution of crossovers along chromosome 3 (sliding window of 2Mb, step size of 50Kb) in female meiosis (RIL 7RV498). Heterozygous regions are represented in purple and homozygous in yellow, and compared to the F1 hybrid (grey). Grey shadings indicate the non/low-recombining zones around the centromeric repeats, and light blue shadings the peri-centromeric regions, as defined in Underwood et al., 2018. The two reciprocal crosses are shown in two different panels and the respective genome structure (the two homologous chromosomes) is shown below each graph (Col-0 regions in blue, Ct-1 regions in red, centromeric region in grey). Transitions between regions of the RILs are shown by dashed grey lines. **C)** Distribution of crossovers in female meiosis in both reciprocal crosses for each of the five *A. thaliana* chromosomes (RIL 7RV498). The 1.17Mb inversion between Col-0 and Ct-1 on the short arm of chromosome 4 is indicated by a purple rectangle. **D)** Heatmaps showing the aggregated CO frequency for each interval in both reciprocal crosses.

Indeed, we could detect 10 times more double-COs on chromosome 3 (Fig. 2A) when polymorphism is present at chromosome ends. Most of these double events are in fact positioned at the extremities on the chromosome (Fig. 2A, top), as predicted. At the genome-wide scale, the number of double COs on all chromosomes was increased 3-fold in bRILs compared to F1 hybrids (0.40 events per plant in RIL backcross, v. 0.13 in F1 hybrids, Fig. 2B and Supplementary Fig. S9A). In male data, while the number of double COs was unchanged, the number of triple COs was significantly increased (0.22 events per plant in RIL backcross, vs. 0.09 in F1 hybrids, Supplementary Fig. S9A). The global CO number per individual was therefore increased by 15.8% in female and 14% male bRIL meiosis compared to F1 hybrid (Table S5 ; Supplementary Fig. S6A and S6B).

If more double- and triple-COs occur, this could mean that CO interference strength could be weaker. However, neither inter-CO distance distribution nor Coefficient of Coincidence (CoC) curves were significantly different in the bRILs compared to the F1 hybrid controls (in males where enough double events could be retrieved, Fig. 2D, Supplementary Table S6, Supplementary Fig. S9B). Interference strength therefore did not appear to be altered by the presence of heterozygous and homozygous regions.

Altogether, these results show that reshaping of the CO landscape through polymorphism allows for more double events in females and triple events in males, increasing the number of total of COs formed by ∼15%. This reshaping does not appear to be the result of a reduction in interference strength. All the data shown here argue toward a major effect of the presence of polymorphism as a powerful recombination driver in *A. thaliana*.

### Reshaping of the crossover landscape through polymorphism and its interplay with interference

In both male and female, centromeric regions, encompassing a strictly Non-Recombining Zone (NRZ) and a Low-Recombining zone (LRZ, Fernandes et al., 2024) are depleted for CO events. Further along the chromosome, the recombination landscape show a large enrichment of COs in the 1-2Mb regions surrounding the centromeres in *A. thaliana* (Jing et al., 2024; Rowan et al., 2019; Salomé et al., 2012), and this has been a persistent feature throughout the species evolutionary history (Lian, Solier, et al., 2022). For simplicity, we will refer to these regions as peri-centromeric High-Recombining Zones (HRZ), and we define them as regions with above genome average DNA methylation rate (as in Underwood et al., 2018). With this definition, the HRZ actually includes the NRZ and LRZ, which contribute very little to the total number of COs in this region. The recombination drivers that promote CO formation in these regions are unknown (see also Discussion).

We studied the recombination frequency within the eight peri-centromeric HRZ on the long arms of the five chromosomes (Supplementary Fig. S3A). In our bRILs, COs tend to form less often in homozygous peri-centromeric HRZ than in F1 hybrids (Fig. 3A and Supplementary Fig. S10, mean of 0.139 CO per interval per plant for F1 hybrid and 0.088 for bRIL homozygous). This depletion in the peri-centromeric HRZ was almost systematically associated with an increase in CO rate in adjacent heterozygous regions further away from the centromere (illustrated in Fig. 3B-D). This suggest that presence of polymorphism in chromosome arms promotes CO formation at the expense of the peri-centromeric regions.

This can be easily explained by CO interference, by which the locations of COs along one chromosome are not independent from one another. In the bRILs, COs tend to be driven towards polymorphic regions, siphoning off all events from the other parts of the chromosome. This suggests that presence of polymorphism is a powerful recombination driver, which can compete with regular drivers that usually favor recombination in peri-centromeric HRZ.

Interestingly, heterozygous peri-centromeric HRZ in the bRILs can also show less COs than the F1 hybrid controls, although this difference is not associated with statistical significance at the genome-wide level (Fig. 3A and Supplementary Fig. S10, mean of 0.139 CO per interval per plant for F1 hybrid and 0.116 for bRIL heterozygous). Again, this can be explained by the global redistribution of COs towards heterozygous regions and its interplay with CO interference, as illustrated in Fig. 3D. Whenever a heterozygous peri-centromeric HRZ recombines less than its F1 hybrid counterpart, another region further on the chromosome (including, or not, the peri-centromeric HRZ itself) recombines more. In other words, given the choice, COs tend to favor heterozygous regions, be it in peri-centromeric HRZ or not. This suggests that presence of polymorphism is a stronger recombination driver than the peri-centromeric drive, and can overhaul the whole CO landscape.

**Fig. 2.**
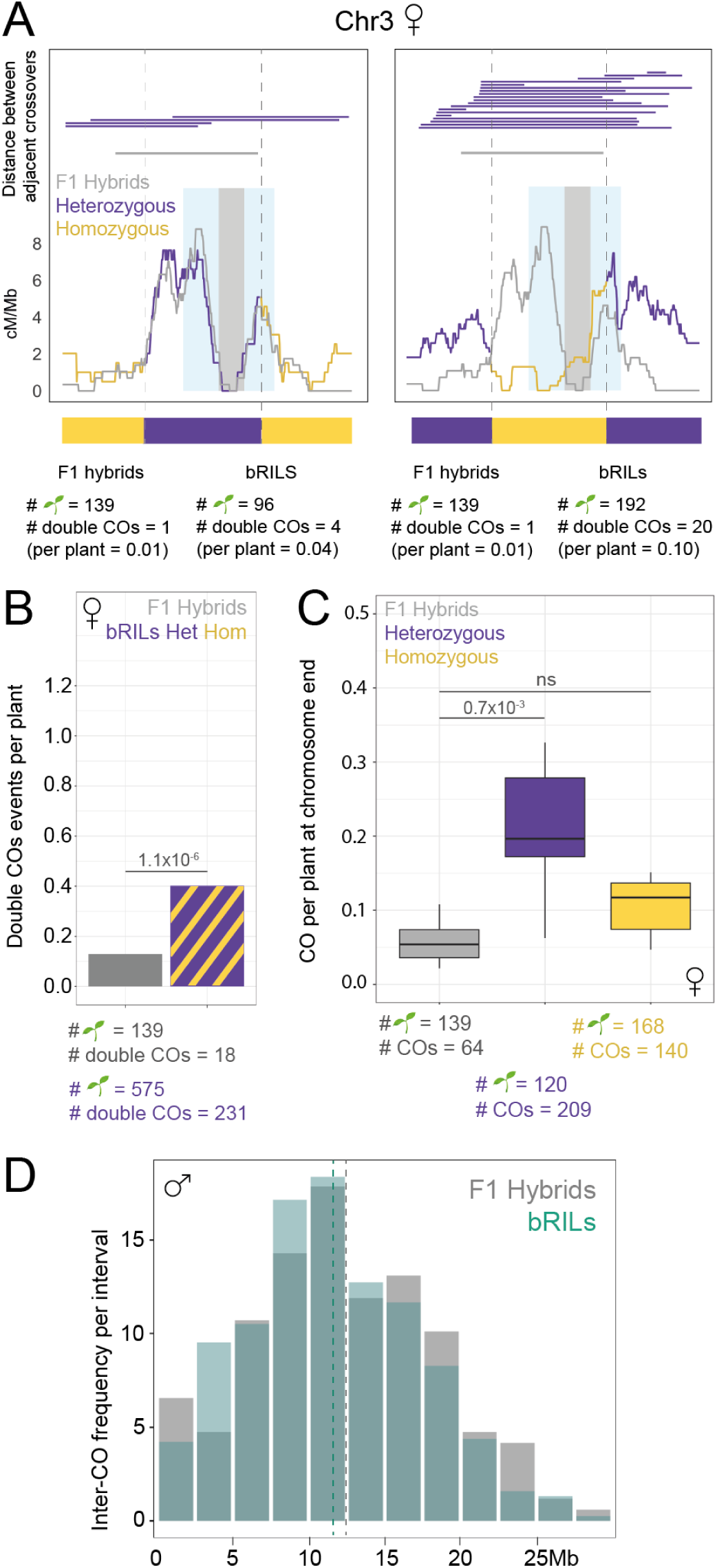
Reshaping of the crossover landscape in presence of polymorphism blocks. **A)** Bottom= Distribution of all COs on Chromosomes 3 (RIL 7RV498) in females meiosis (cM/Mb). Top: Distance and position between adjacent double COs (on the same chromatid). Double COs from bRILs data are plotted in purple and control F1 hybrid events in grey. **B)** The number of double adjacent CO events in the control F1 hybrid populations (grey) and female bRIL populations (striped purple and yellow). Chi-square was used to calculate statistical significance. **C)** Analysis of the mean number of COs formed per plant at eight chromosome ends in female bRILs and F1 hybrids. Kruskal-Wallis test was used to calculate statistical significance. **D)** Inter-CO distance between adjacent double and triple CO events in male bRIL and F1 hybrids populations. The dashed lines represent the mean distances in both populations.

**Fig. 3.**
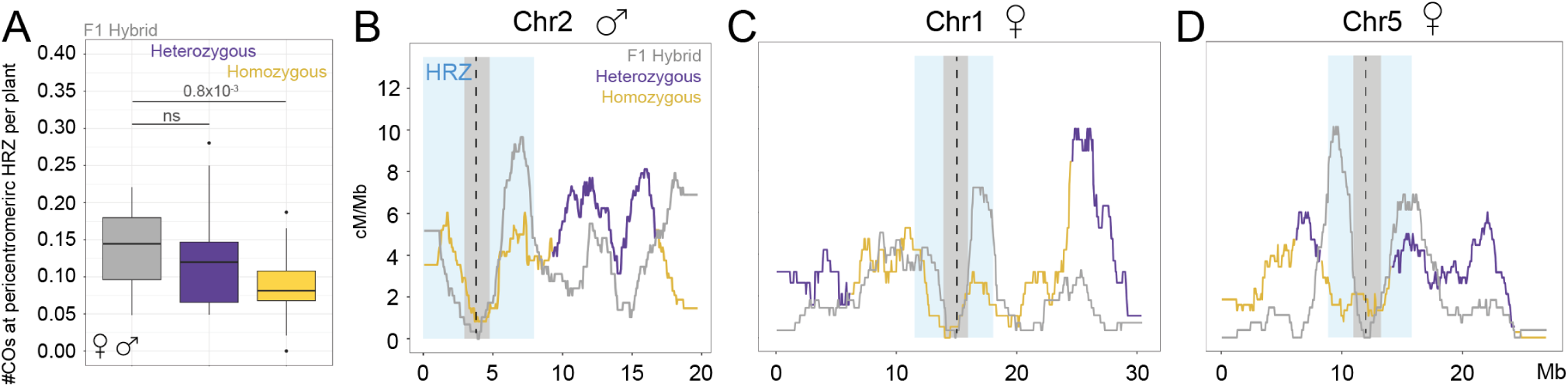
Delocalization of crossovers from peri-centromeric HRZ in presence of polymorphism blocks. **A)** Analysis of the mean number of COs formed per plant within peri-centromeric HRZ, as defined by Underwood et al., 2018, which include the NRZ and the LRZ (in grey, Fernandes et al., 2024) See also Supplementary Fig. S3A. Kruskal-Wallis test was used to calculate statistical significance. **B)** Distributions of COs at chromosome 2 in male bRILs. **C)** Distributions of COs at chromosome 1 in female bRILs. **D)** Distributions of COs at chromosome 5 in female bRILs.

### MSH2 is required for the positive effect of polymorphism on crossover rate

How can the presence of polymorphism promote CO formation? The DNA mismatch repair (MMR) is a conserved machinery that can recognize, and repair, DNA mismatches (be it mis-matched nucleotides or presence of small insertions/deletions). The conserved MutS complex, composed of an heteroduplex between Msh2 and either Msh3 or Msh6, can bind to a mismatch and recruit downstream partners to remove and replace the aberrant nucleotides (reviewed in Manhart and Alani, 2016). During meiotic CO formation, the DNA molecule that receives a DSB invades a homologous template to initiate the repair. If the homologous chromosome bears SNPs or small INDELs compared to the broken molecule, mismatches can occur through the formation of heteroduplexes, at the invasion step or later in the processing of recombination intermediates. We therefore tested whether the MMR could participate in favoring CO formation in polymorphic regions, with the hypothesis that MSH2 could detect mismatches created during DSB repair.

We generated full knock-out mutants of *msh2* using CRISPR/Cas9 in all appropriate lines, and performed crosses as in Supplementary Figure S1. Two *msh2 -/-* bRIL populations of 152 (RIL 7RV498 backcrosses to *msh2 -/-* Col-0) and 136 (backcrossed to *msh2 -/-* Ct-1) individuals were sequenced. Among these 288 individuals, no sequencing reads mapped to the *msh2* coding sequence, confirming the mutant status of all the sequenced plants (Supplementary Fig. S11).

In these *msh2 -/-* bRILs, heterozygous regions regions recombine less than in wild-type bRILs (Fig. 4A, 0.043 CO/Mb per interval per plant in wild-type bRILs and 0.033 in *msh2* bRILs), and conversely, homozygous regions recombine more (0.032 CO/Mb per interval per plant in wild-type and 0.041 in *msh2*). This suggests that in absence of MSH2, COs are no longer redistributed to polymorphic regions as observed in wild type, and that MMR is required to drive recombination to polymorphic regions. This is further illustrated by the fact that the recombination rate is either similar and most often elevated in homozygous regions (yellow) in *msh2* compared to wild-type bRILs (Fig. 4D and E). Reversely, the recombination rate is either similar and most often lower in heterozygous regions (purple). This is consistent with previous studies showing that MSH2 was necessary for the juxtaposition effect (Blackwell et al., 2020; Dluzewska et al., 2023).

**Fig. 4.**
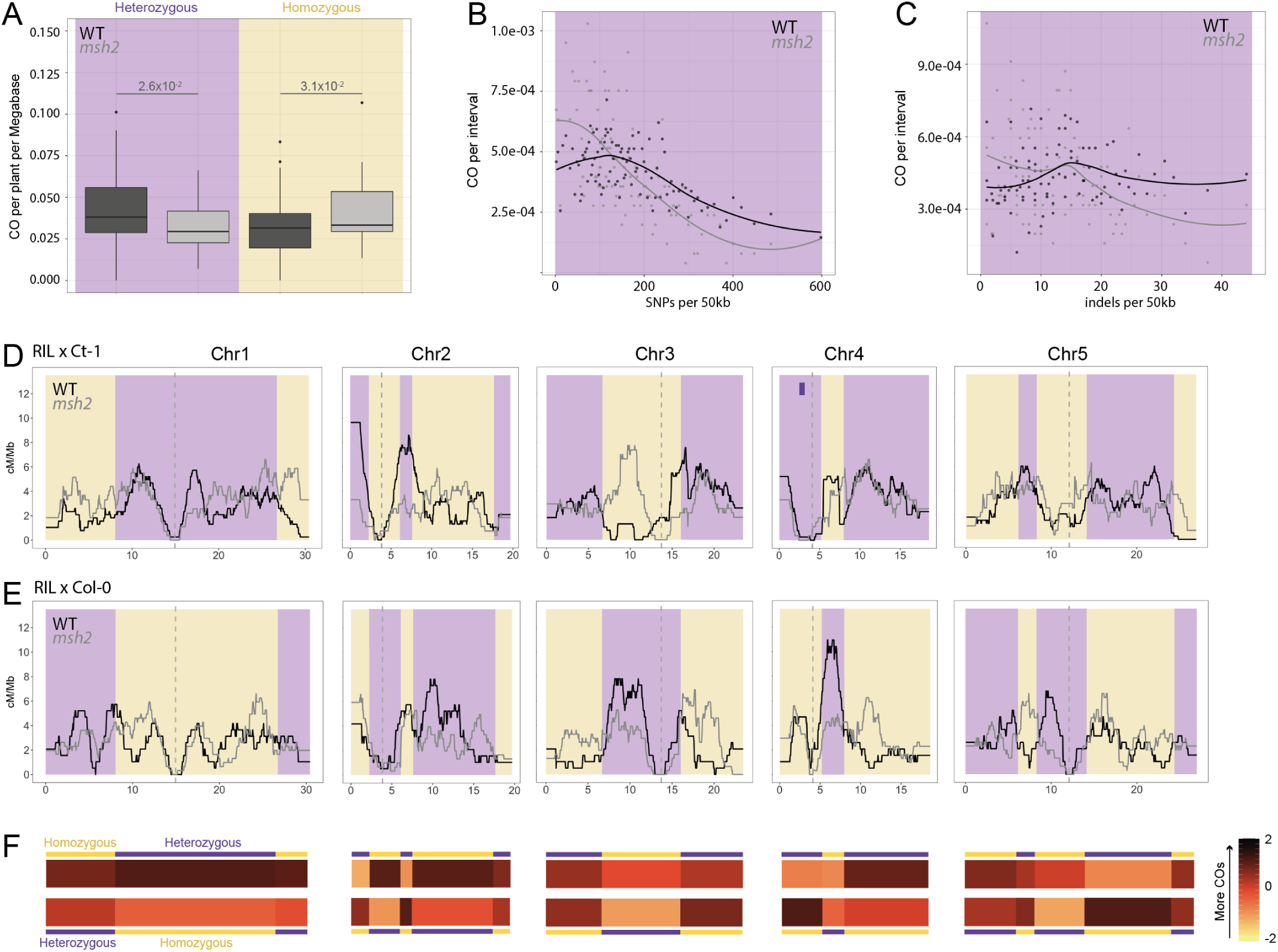
In absence of MSH2, COs tend to form in non-polymorphic regions of the genome. **A)** Normalized number of crossovers per plant in female and male meiosis in WT bRILs (dark grey) and *msh2* bRILs (light grey) (yellow and purple, for homozygous and heterozygous regions respectively). Mann-Whitney test was used to calculate statistical significance. **B)** 50kb windows across heterozygous regions were scored for SNP densities, and the fraction of normalized crossovers compared to per 50kb in WT and *msh2* bRILs. **C)** 50kb windows across heterozygous regions were scored for indel densities, and the fraction of normalized crossovers compared to per 50kb in WT and *msh2* bRILs. **D-E)** Distribution of crossovers in female meiosis in both reciprocal crosses for each of the five *A. thaliana* chromosomes (sliding window of 2Mb, step size of 50Kb) in WT F1 hybrid and *msh2* bRILs. The 1.17Mb inversion between Col-0 and Ct-1 on the short arm of chromosome 4 is indicated by a purple rectangle. Dashed lines indicate the center of the centromeric repeats. **F.** Heatmaps of the CO frequency within the same intervals in both reciprocal crosses.

We then looked at individual 50kb windows within heterozygous regions across the genome. Because polymorphism density is not homogeneous across the genome, windows display between 0 to more than 500 SNPs per 50kb. When looking at wild-type bRILs, COs occurring in heterozygous regions are only weakly affected by the level of polymorphism, be it SNPs or indels, within the heterozygous regions (black, Fig. 4B and C). However in *msh2 -/-* bRILs, windows with the fewer SNPs and indels displayed the most CO events, showing that MSH2 is required for COs to form in polymorphic intervals within heterozygous regions.

This is recapitulated when looking at the total number of crossovers per interval (Fig. 4F), where homozygous regions display more COs (darker color palette) than heterozygous ones (less COs, paler colors). This is true for both female and male (Fig. 4 and Supplementary Fig. S12). This is the opposite image of what is observed in wild-type bRILs, showing that MSH2 is required to re-distribute COs to polymorphic regions of the genome.

To conclude, the absence of MSH2 leads to COs forming preferentially in homozygous regions when given the choice. In turns, this points to a pro-CO role of MMR in *A. thaliana*, and that in absence of the MMR, the homologous recombination machinery tends to promote CO formation in non-polymorphic regions.

### Crossovers are formed homogeneously along heterozygous intervals

We looked at the position of COs within 1Mb (Fig. 5A) and 2 Mb intervals (Fig. 5A) on each side of multiple frontiers between polymorphic and non-polymorphic regions (Supplementary Fig. S3B). Nineteen intervals of 1Mb and thirteen intervals of 2Mb on each side of the frontiers were selected (see Methods) and the number of events was pooled in 0.5 Mb windows. Across all frontiers, we observed a sharp transition between homozygous and heterozygous regions (Fig. 5A and B), recapitulating our previous results that more COs are formed in heterozygous regions than homozygous ones. Additionally, this increase appeared to be similar throughout once the frontier was reached, with no significant differences across the different 0.5Mb windows. The same trend was observed in both female and male data independently (Supplementary Fig. S13).

**Fig. 5.**
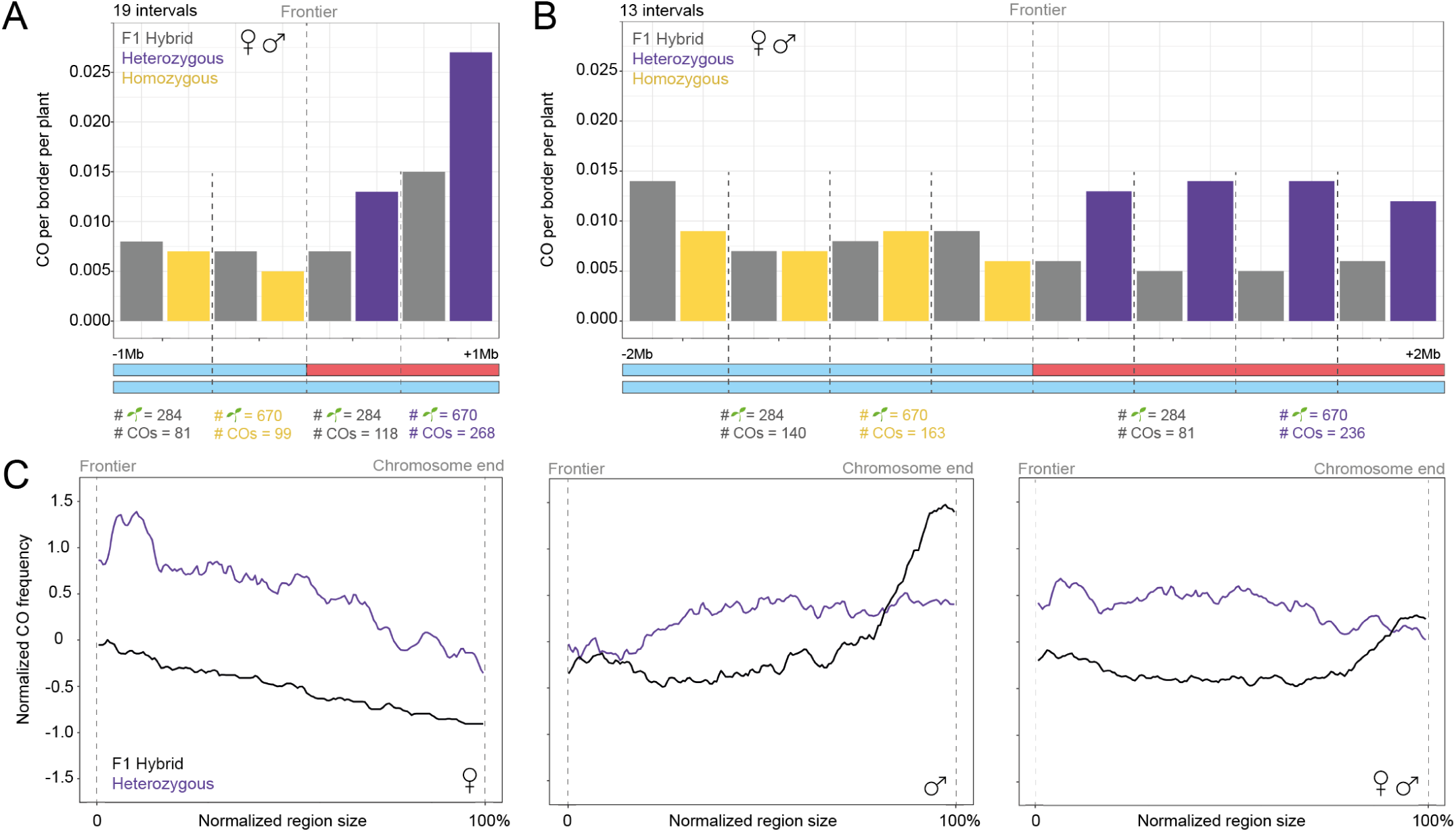
Description of the crossover landscape at frontiers between polymorphic and non-polymorphic regions and within heterozygous intervals. **A)** The number of COs across 19 frontiers (Supplementary Fig. S3B) within the two RILs, male and female data aggregated. The position of the COs was retrieved within 1Mb around the frontiers, and the numbers of events were plotted in four 0.5Mb windows. **B)** The same analysis was done for 13 intervals of 2Mb around the frontiers, yielding eight windows of 0.5Mb (see Methods and Supplementary Fig. S3)B). This analysis was done using only the Col-0 EMS mutations markers for all intervals (both homozygous and heterozygous), to avoid differences in resolution. **C)** The CO frequency normalized within 7 intervals at chromosome ends (Supplementary Fig. S3C, sliding window of 2Mb, step-size 50Kb) was normalized in two-hundreds windows and then aggregated for (i) female, (ii) male, (iii) female and male. This analysis was done using only sequence Col-0/Ct-1 divergence as markers.

To go further, the distributions within whole heterozygous intervals were aggregated in order to detect potential patterns. Seven intervals of similar sizes (between 5.8Mb and 8Mb, see Methods and Supplementary Fig. S3C) and located at similar positions along chromosomes (sub-telomeric) were pooled to observe general trends. Because the intervals are at chromosome ends, the recombination frequency in the F1 hybrids reflects the male/female differences observed previously, with high frequency of COs in males at the distal ends and fewer events in females. Confirming our previous observations, more COs were recorded in all these heterozygous regions in the bRILs than in the F1 controls, except in the distal part of chromosomes in male meiosis (Fig. 5C), corroborating the fact that the CO landscape is globally reshaped in our lines. In female meiosis, there was an increase within the first 15% (∼1Mb) at the frontier (Fig. 5C, female panel). For males however, no such effect was observed, with the same number of COs formed compared with F1 hybrid controls close to the border (Fig. 5C, male panel). Finally, after aggregation of male and female data, no specific pattern could be observed, with a flat trend all along the heterozygous interval compared to F1 hybrids (Fig. 5C, right panel).

A previous study proposed a model which showed a skewed distribution close to the frontier between a homozygous and heterozygous region, with more events being formed within the first 1-2Mb after the border (Dluzewska et al., 2023). This effect was measured with recombination markers expressed in seeds, aggregating both female and male meiosis effects. Here, we describe extensively how the effect acts along heterozygous intervals all across the genome, independently in both sexes. Our global analyses of multiple intervals across all five chromosomes of *A. thaliana* failed to highlight a specific pattern in CO distribution within heterozygous regions, leading to the hypothesis that polymorphism would favor CO formation uniformly along heterozygous intervals.

## Discussion

What mechanisms regulate CO distribution along chromosomes? In this study we aimed at understanding the impact of polymorphism on this distribution by leveraging the genetic structure of Recombinant Inbred Lines in *A. thaliana*. These lines allow us to assess the recombination landscape in a genetic context with altered polymorphism distribution compared to highly homozygous, evolved, wild lines.

We uncovered that presence of polymorphism is a powerful driver for recombination, competing with other recombination drivers that have shaped the species conserved CO landscape. In many eukaryotes, profile of recombination along chromosomes and polymorphism density are closely associated (Andersen et al., 2012; Begun and Aquadro, 1992; Burri et al., 2015; Dvorřák et al., 1998; Lian, Solier, et al., 2022; Nachman, 2001; Stephan and Langley, 1998; Tenaillon et al., 2001). This is thought to arise mainly because selection does not only select for alleles, but for whole haplotypes (chromosomal segments between two adjacent recombination events, Ellegren and Galtier, 2016). This means that when the recombination rate is low, many neutral polymorphisms remain linked to a selected allele. When this allele is selected for (or against) in a population, the whole haplotype becomes prevalent (or disappears, respectively), leading to a decrease in diversity in the region. Conversely, when recombination rate is high, selected haplotypes are shorter in length, and more neutral polymorphisms around the selected allele can persist in the population. This positive association between recombination rate and sequence divergence can be reinforced by the slight mutagenic effect of recombination, in the order of ∼1 SNP per million cells in fission yeast, and 1 in 4 sperms in human (Arbel Eden and Simchen, 2019; Duret and Arndt, 2008; Hinch et al., 2023).

However, mechanistically, polymorphism is a known barrier to recombination. In non-meiotic cells, as little as 0.3-0.5% sequence divergence (1 SNP every 300-500bp) drastically reduces recombination in *Saccharomyces cerevisiae* and *A. thaliana* (Datta et al., 1997; L. Li et al., 2006). During meiosis, COs also appear to avoid interhomolog polymorphism, recombining mostly with identical sequences in yeast (Alani et al., 1994; Borts and Haber, 1987; Chambers et al., 1996; Raffoux et al., 2018). In *A. thaliana* and *S. cerevisiae*, the study of recombination hotspots demonstrated a trend for COs to be formed in less polymorphic loci (Cooper et al., 2018; Serra et al., 2018; Szymanska-Lejman et al., 2023). Beyond this local effect, on the chromosome scale, previous data indicated that the presence of polymorphism at the end of chromosome 3, when the rest of the chromosome did not present any sequence differences, could boost recombination in the polymorphic interval (Blackwell et al., 2020; Dluzewska et al., 2023; Ziolkowski et al., 2015). In this study, we show that presence of polymorphism can change the relative recombination rate throughout the genome: globally, all heterozygous regions recombine more than homozygous ones, independently of their genomic location (Fig. 1).

A highly conserved feature of the recombination landscape is the centromeric inhibition of CO formation (Fernandes et al., 2019). This repression would be essential for the correct segregation of homologs during meiosis I, where COs could interfere with proper cohesin maintenance at centromeres and microtubule attachment (Zielinska et al., 2019). The regions flanking the centromeric repeats are typically devoid of COs in many species such as maize, tomato, or male stickleback fish (Ganal et al., 2011; Sim et al., 2012; Venu et al., 2024). Nonetheless, a very striking characteristic of *A. thaliana* CO distribution is a the presence of High-Recombining Zones (HRZ) flanking the centromeric region. In males, around a third of all events occur in these regions, when they account for only 20% of the genome. In females, the effect is even more drastic, with half of all COs formed in peri-centromeric HRZ. The underlying drivers of this localization are not known (see also below). However, our results show that these regions can be largely depleted in COs if polymorphic regions are located further on chromosomal arms (Fig.2). This polymorphism-based recombination driver is strong enough not only to deplete peri-centromeric HRZ of COs, but also to promote recombination in the distal parts of chromosome arms in female, where very few COs are usually observed. Therefore, we demonstrate that the presence of polymorphic and non-polymorphic regions within the same homologous pair can massively meddle with what was previously thought as inescapable patterns of *A. thaliana* CO distribution.

Contrary to what was previously reported for chromosome 3 (Dluzewska et al., 2023), we were unable to detect an enrichment for COs at the frontiers between polymorphic and non-polymorphic regions, and we show that the profile of recombination across heterozygous intervals is homogeneous (Fig.3). On chromosome 3 itself, where a frontier is present at a similar location as in Dluzewska et al., 2023, we could observe an increase close to the frontier in female data, but not in males (Supplementary Fig.S7 & S8). This could be due to the fact that we do not detect enough events (236 COs on a 2Mb interval, Fig.3; *vs.* ∼4,000 in Dluzewska et al., 2023). However, we analyzed many borders across the genome and we believe that the fact that we could not detect any pattern along 7 intervals on 3 chromosomes (Fig.3) suggests that this border effect, if it exists, is small.

Overall, our observations of the interplay between polymorphism and CO interference leads us to propose a model (Fig. 6A) where the choice for a DSB (green dots) to become a CO is influenced by peri-centromeric recombination drivers in F1 and pure lines, heavily skewing the CO distribution towards regions surrounding the centromere (black profile). In genetic backgrounds where the polymorphism density is uncoupled from the historical recombination pattern, as in our bRILs, the presence of polymorphism becomes a powerful recombination driver (purple and yellow profile), superseding the peri-centromeric drivers. Together with interference (*d_int_*), this polymorphism driver drastically alters the chromosome-wide recombination profile.

**Fig. 6.**
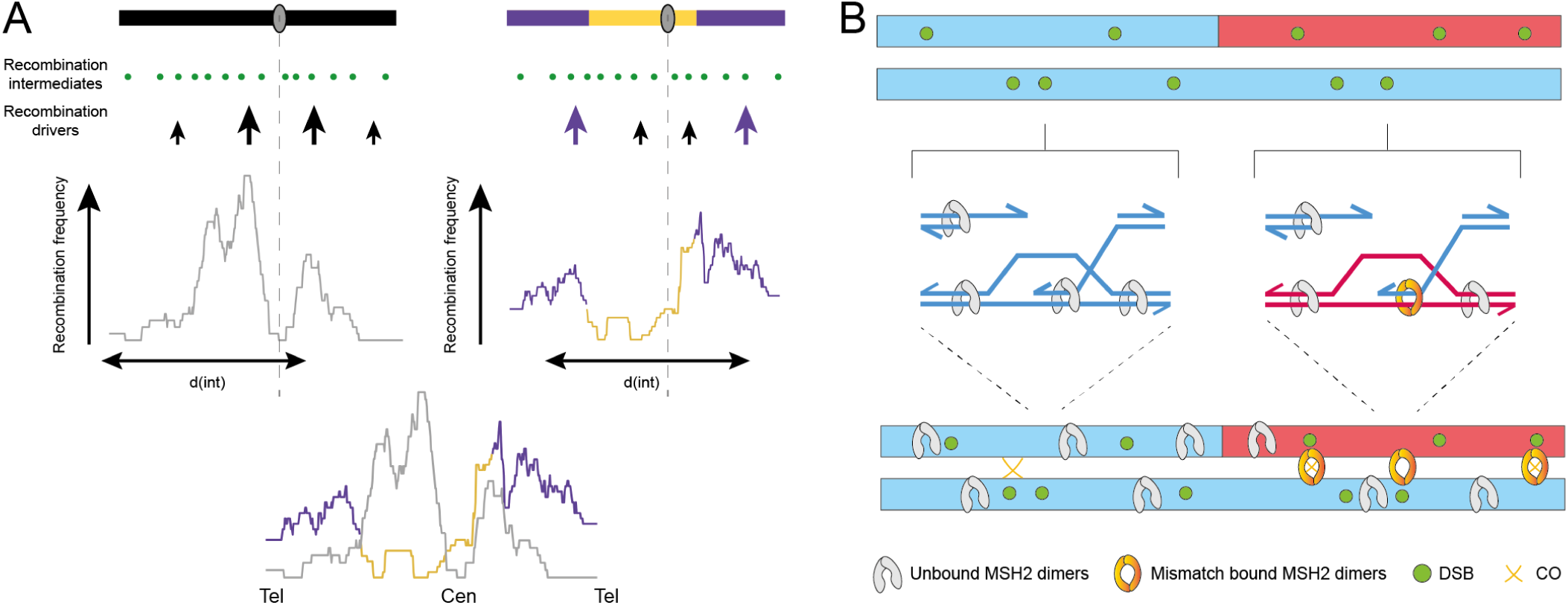
Model of polymorphism as a major recombination driver in *A. thaliana*. **A)** In fully homozygous or heterozygous backgrounds (black), the CO landscape is mainly influenced by peri-centromeric recombination drivers. However, when a polymorphic region is present at chromosome ends, presence of polymorphism favors recombination in polymorphic regions (purple). Interference also heavily impacts the landscape, and while the distance between CO events d(int) is unchanged, the relocation of COs away from the center of the chromosome allows for more events to be formed. **B)** The mismatch repair (MMR) protein MSH2 would be distributed evenly all along chromosomes with weak affinity for DNA. During recombination, mismatches can occur in heterozygous regions within heteroduplexes, which would recruit and MSH2 and promote an ATP-driven conformational change, in turn favoring the formation of COs in polymorphic regions.

Mechanistically, what are these recombination drivers? Regarding the peri-centromeric recombination driver, an interesting hypothesis can be drawn from the global recombination maps drawn in hyper-recombinant mutants where interference is abolished (Dluzewska et al., 2023; Jing et al., 2025). In these mutants, COs are not sensitive to interference and the recombination landscape shows no relative enrichment in COs in the regions defined as HRZ in wild type. These are the first mutant backgrounds in which the recombination landscape shows little recombination around the centromeres. This suggests that CO interference itself could be responsible for the high recombination rate in the HRZ around the centromeres in *A. thaliana*.

As for the polymorphism, in *A. thaliana*, we show, in accordance with previous work (Blackwell et al., 2020; Dluzewska et al., 2023; Szymanska-Lejman et al., 2023), that the mismatch repair machinery, through its main effector Msh2, is required to relocate COs to polymorphic regions. This is in contrast to the situation in yeast, where Msh2 restricts CO formation in polymorphic hotspots (Cooper et al., 2018; Martini et al., 2011). These two observations can be reconciled when looking at the right scale: even in *A. thaliana*, recombination tends to be favored in non-polymorphic sequences at the kb-scale (Serra et al., 2018; Szymanska-Lejman et al., 2023). It is therefore most likely that, while on a global scale polymorphism favors CO formation, on a local scale (hundreds of base pairs), polymorphism still interferes with CO formation. We further speculate that in wild-type hybrids, where high levels of divergence are present in peri-centromeric regions, MMR would further increase the propensity of COs to form close to centromeres in *A. thaliana*.

We therefore propose a model in which MSH2 would be loaded all along chromosomes (as observed cytologically, Blackwell et al., 2020) but only capable of recruiting partners upon recognition of a mismatch contained within recombination intermediates (Fig. 6B). It has indeed been proposed that in the absence of mismatches, the MSH complex would show a weak affinity for DNA but upon mismatch binding, the dimer would change conformation (in an ATP-dependent manner), enter a sliding clamp mode and recruit downstream repair factors (Manhart and Alani, 2016). Our hypothesis is that, after binding mismatches within heteroduplexes, Msh2 could recruit pro-COs factors, leading to preferentially form COs. Whether Msh2 can then directly recruit the homologous recombination machinery to promote COs (*e.g.,*, MLH1/3, Rogacheva et al., 2014), or other factors that contribute to favoring CO formation in the region, remains to be tested. Although we cannot exclude that DSB distribution could be shifted in the bRILs, we do not have any hypothesis about how it would be possible to detect the presence of polymorphism before the two homologous chromosomes come into close proximity, which occurs after DSB formation.

Why would *A. thaliana* recombine in polymorphic regions? Like many plants, it is a self-pollinated species and its populations tend to be homozygous (Lande and Schemske, 1985; Stebbins, 1957; Wright et al., 2013). Recombining in heterozygous regions can slow down the loss of heterozygosity by reducing the size of haplotypes. On the other hand, many plants are polyploids who need to avoid recombination between non-homologous sequences to only form COs between homologous and not between hom**e**ologous chromosomes, to ensure faithful segregation of chromosomes during meiosis. In wheat, the plant specific mismatch repair protein MSH7 appears to be crucial for the distinction between homologs and homeologs (Serra et al., 2021). This suggests that there could be a threshold in divergence levels at which the MMR switches from pro- to anti-CO. It would be very interesting to test whether polymorphism, and Msh2, have the same effect in closely related *Arabidopsis* polyploid species such as *A. suecica*. Another question is whether this effect would be observed with higher degree of divergence between homologs. As seen in Fig. 4B, higher level of polymorphism (300 SNPs/50kb) seem to deter CO formation, in favour of mildly divergent (150 SNP/50kb) intervals.

In conclusion, our work demonstrates that polymorphism is a major actor of the CO landscape in *A. thaliana*, and suggests that polymorphism can be a powerful tool for (i) breeders to reduce linkage drag to facilitate the introgression of new traits into elite genomes; (ii) studying fundamental aspects underlying CO formation and designation.

## Data Availability

All sequencing data is available from ArrayExpress: 7RV168 and 7RV498 deep sequencing (E-MTAB-15108), and the different CO mapping populations will be soon submitted to the same repository.

## Contributions

CG conceptualized the project. BM and CG produced the data. LLM and SP helped with generating the plant material. LQ, SM, YM and BM wrote the code to detect crossovers. BM analyzed the data and made all the figures. BM and CG wrote the paper.

## Acknowledgments

We thank: The Imagerie-Gif Facility for access to the microscopy facility, supported by l’Agence Nationale de la Recherche (FBI ANR-24-INBS-0005 BIOGEN, SPS ANR-17-EUR-0007 and EUR SPS-GSR); Bruno Huettel and his team from the Max Plank Genome Center (Cologne) for DNA extraction, library preparation and sequencing; Claire Toffano-Nioche for her help with SnakeMake; Raphaël Mercier for the EMS lines; Ian Henderson for his support and R code; Bertrand Llorente, Christine Mézard, Pierre Grognet, Julie Clément, Eric Jenczewski, Eric Espagne and Denise Zickler, for feedback; Aurélie Chambon for her help with cytology sample preparation; Mathilde Grelon for the antibodies and her mentorship; Lionel Gissot for the GoldenBraid plasmids; Hernan Lopez for counseling on CRISPR-Cas9; Veronique Couvreux from the greenhouse; The Versailles Arabidopsis Stock Center (VASC) and Christine Camillieri for the RILs; Laurence Cromer and Mariana Tiscareno Andrade for their help with plant transformation; Nelly Sainsard and Cristina Delwarde from I2BC kitchen; The I2BC’s IT support team (SICS) for making the high performance computing cluster accessible at the I2BC.

## Supplemental Information

**Fig. S1.**
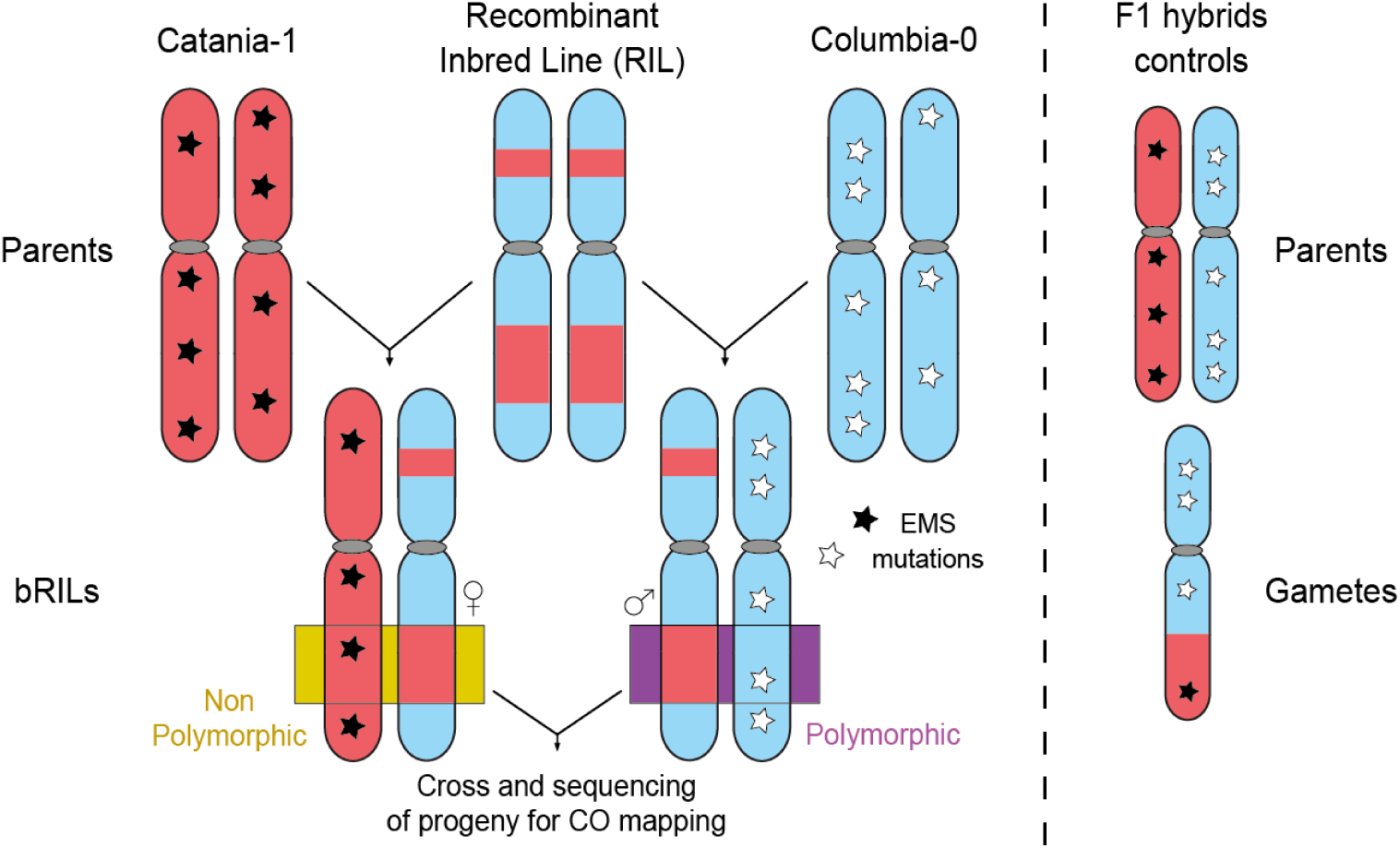
Crossing plan between a Recombinant Inbred Line (RIL) and its parents (Catania-1 and Columbia-0). The backcross between a RIL and its parents lines treated with EMS yields F1 lines, henceforth called bRILs, within which polymorphic and non-polymorphic regions are present. Studying both reciprocal backcrosses allows for the study of every region of the genome under their polymorphic and non-polymorphic state. A cross between two bRILs was made to generate the population that was sequenced in order to detect COs that occurred in the bRIL female and male meioses. F1 hybrids controls were generated by a cross between the same EMS treated parents.

**Fig. S2.**
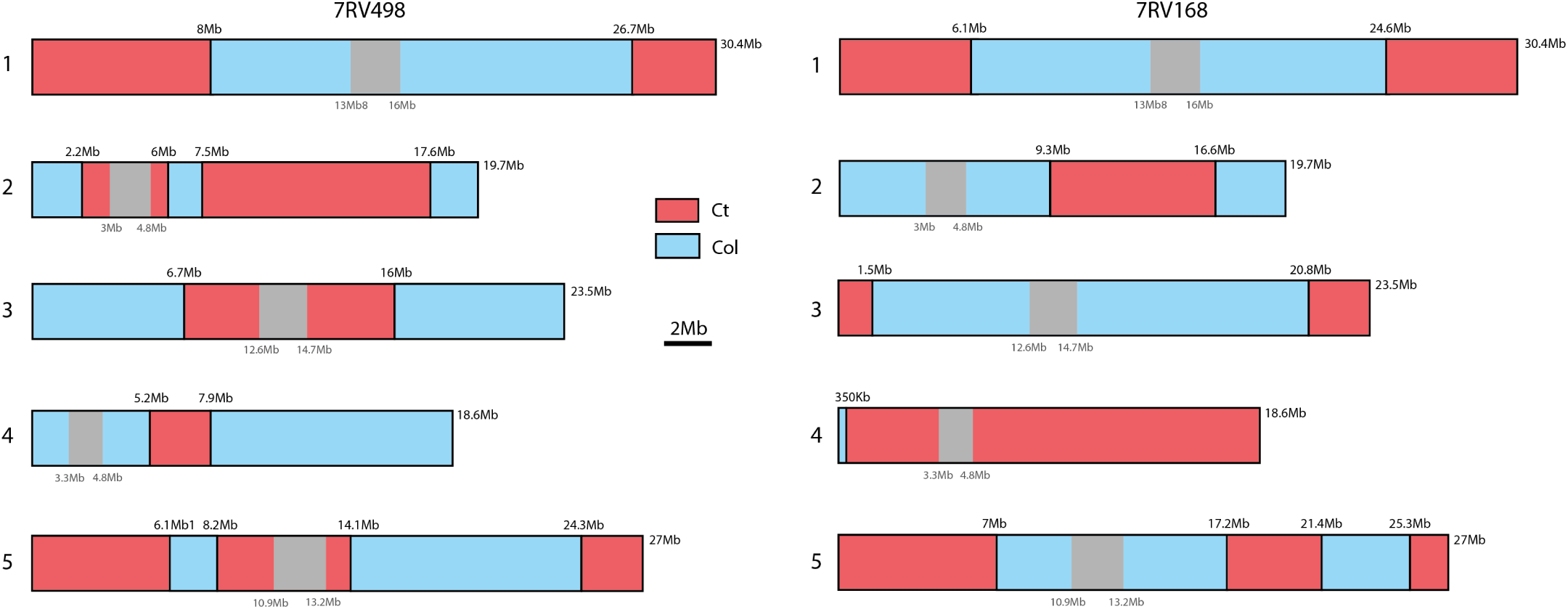
Recombinant Lines 7RV498 and 7R168. These RILs were retrieved from the *A. thaliana* stock center localized in Versailles. The parents were Col-0 (in blue) and Ct-1 (in red). Every chromosome present at least one transition between the two parental genomes. Centromeres are shown in grey.

**Fig. S3.**
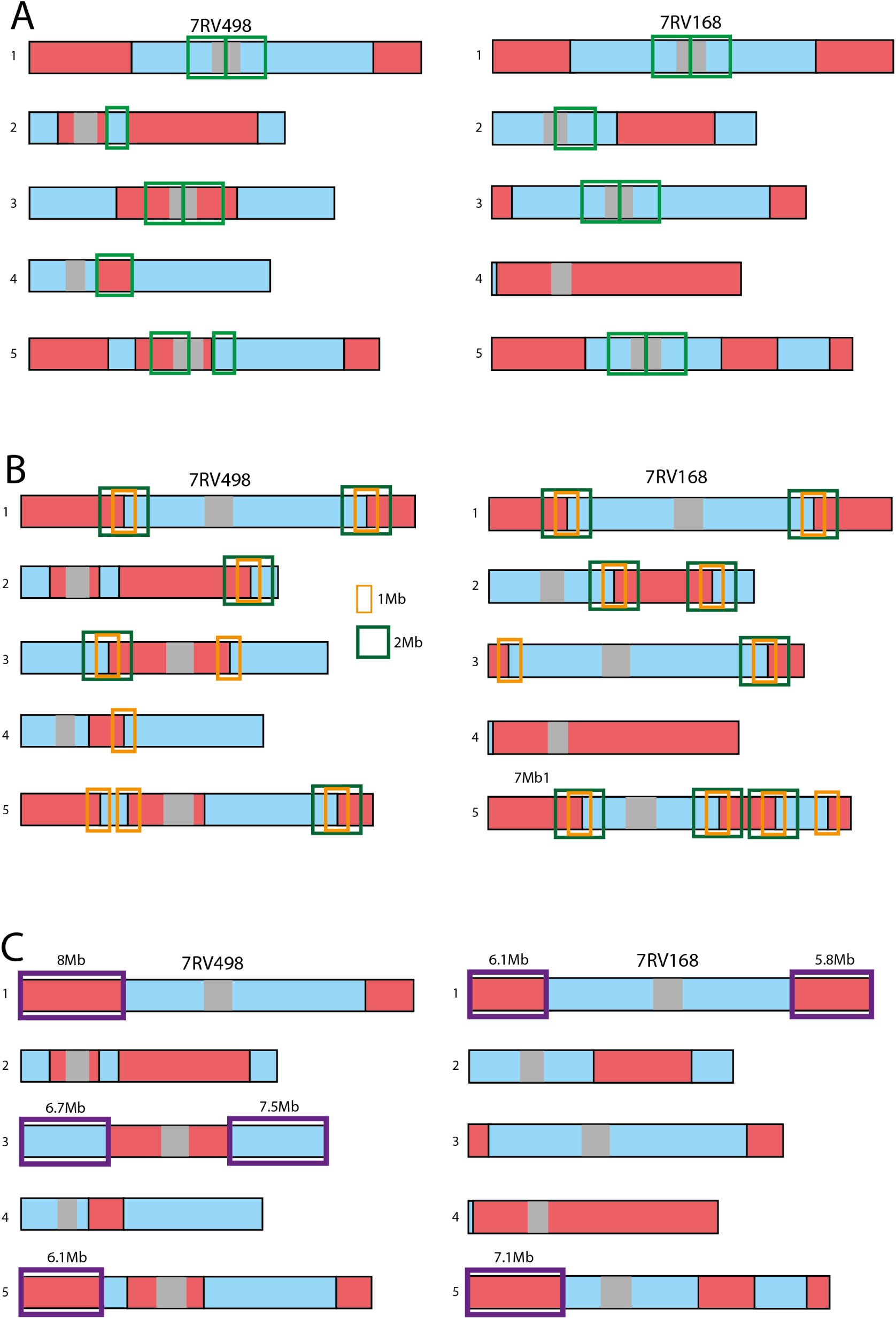
Intervals retrieved for secondary analyzes. A. Pericentromeric regions. B. 1-2Mb intervals around frontiers. C. Chromosome ends intervals for aggregation of heterozygous intervals.

**Fig. S4.**
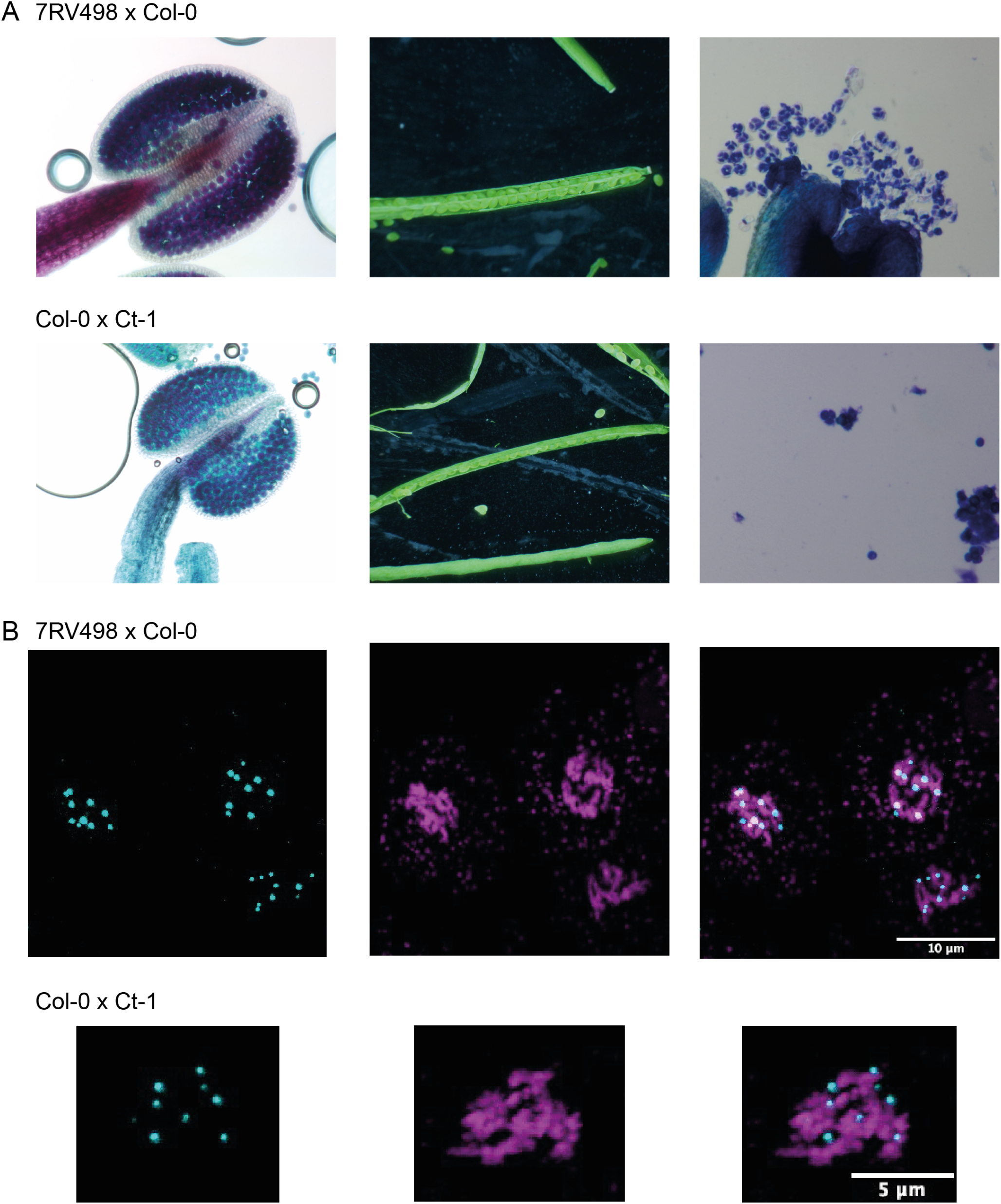
No meiosis defect in F1 hybrids and RIL backcrosses. **A)** Alexander staining of anthers, siliques and toluidine blue staining of tetrads. **B)** Immunolocalization of MLH1 foci was performed: DNA can be observed in magenta (DAPI staining) and MLH1 foci in cyan (anti-MLH1 antibody).

**Fig. S5.**
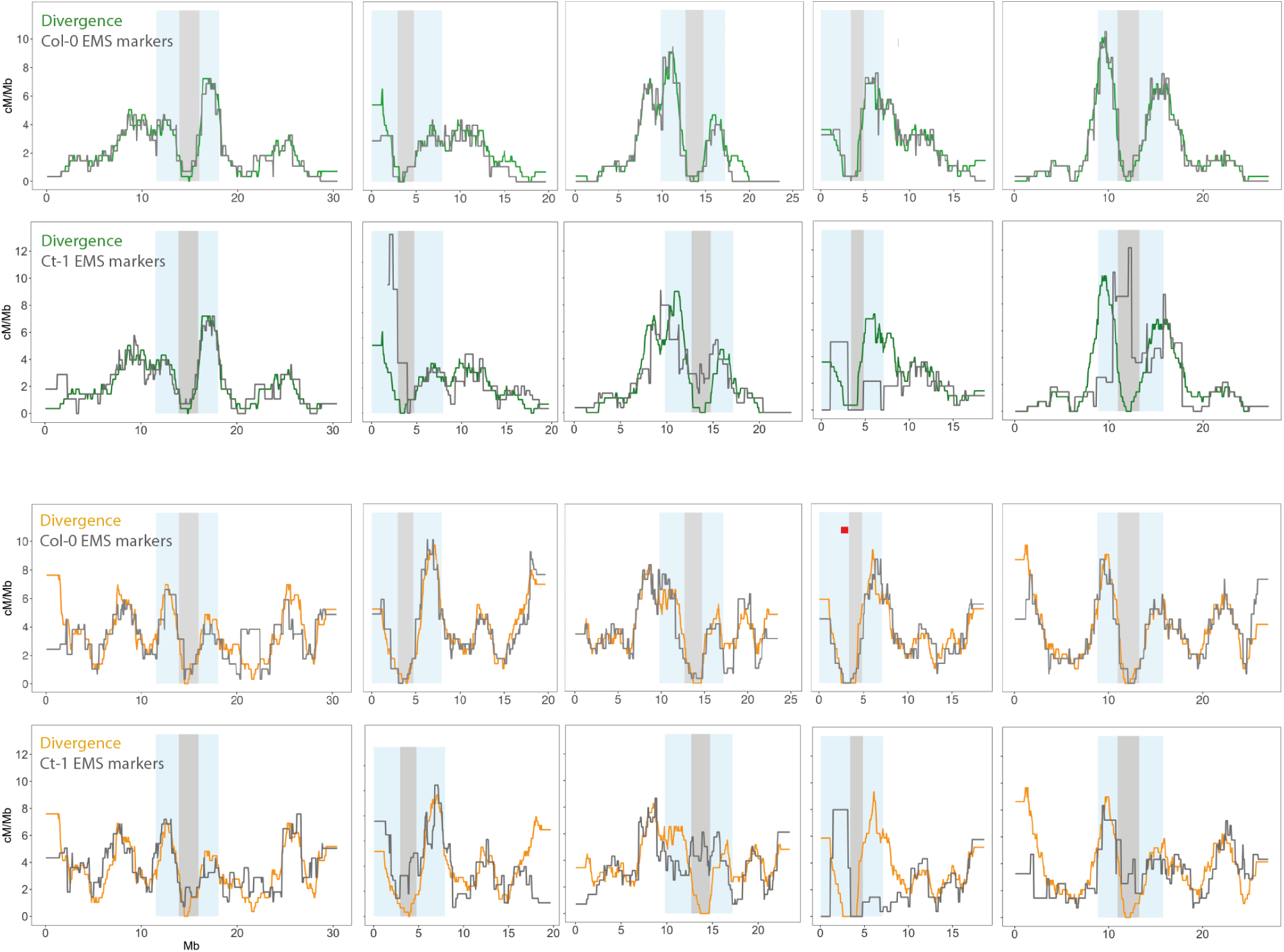
Comparison of distributions depending on the sets of amrkers used for CO detection.. Three sets of markers were used: the Col-0 / Ct-1 divergence, the Col-0 EMS markers and the Ct-1 EMS markers. Female population on the two top panels, male population on the two bottom panels.

**Fig. S6.**
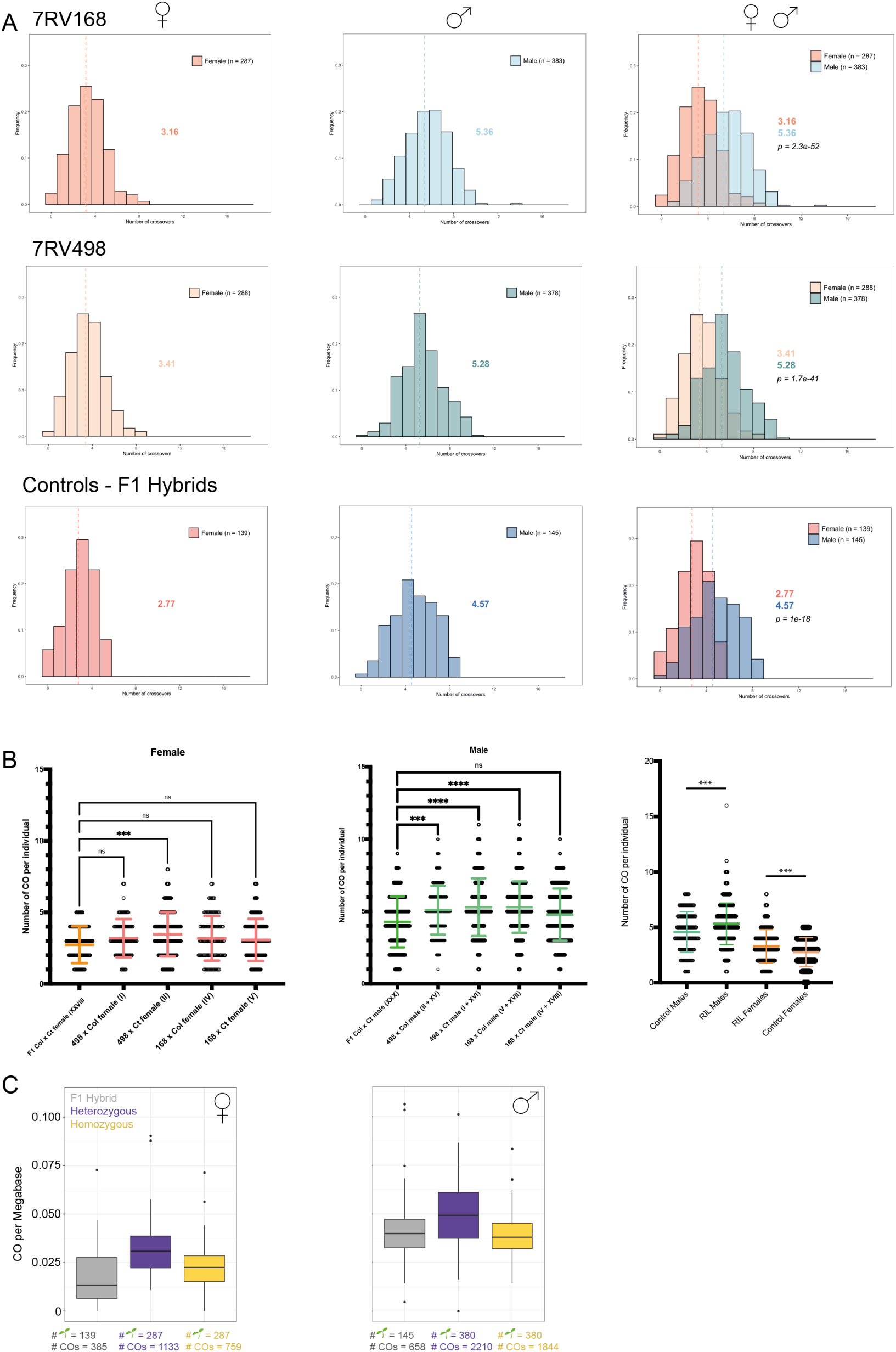
Distribution of the number of crossover per individuals. **A)** For each RIL (7RV168 and 7RV498), both backcrosses populations were pooled. Distribution of the number of crossover per individuals. **B)** For each RIL (7RV168 and 7RV498), each backcross population was analyzed singly and compared to the F1 hybrid control with the Kruskal-Wallis test. **C)** The normalized number of crossovers within 35 intervals per plant in female and male meiosis.

**Fig. S7.**
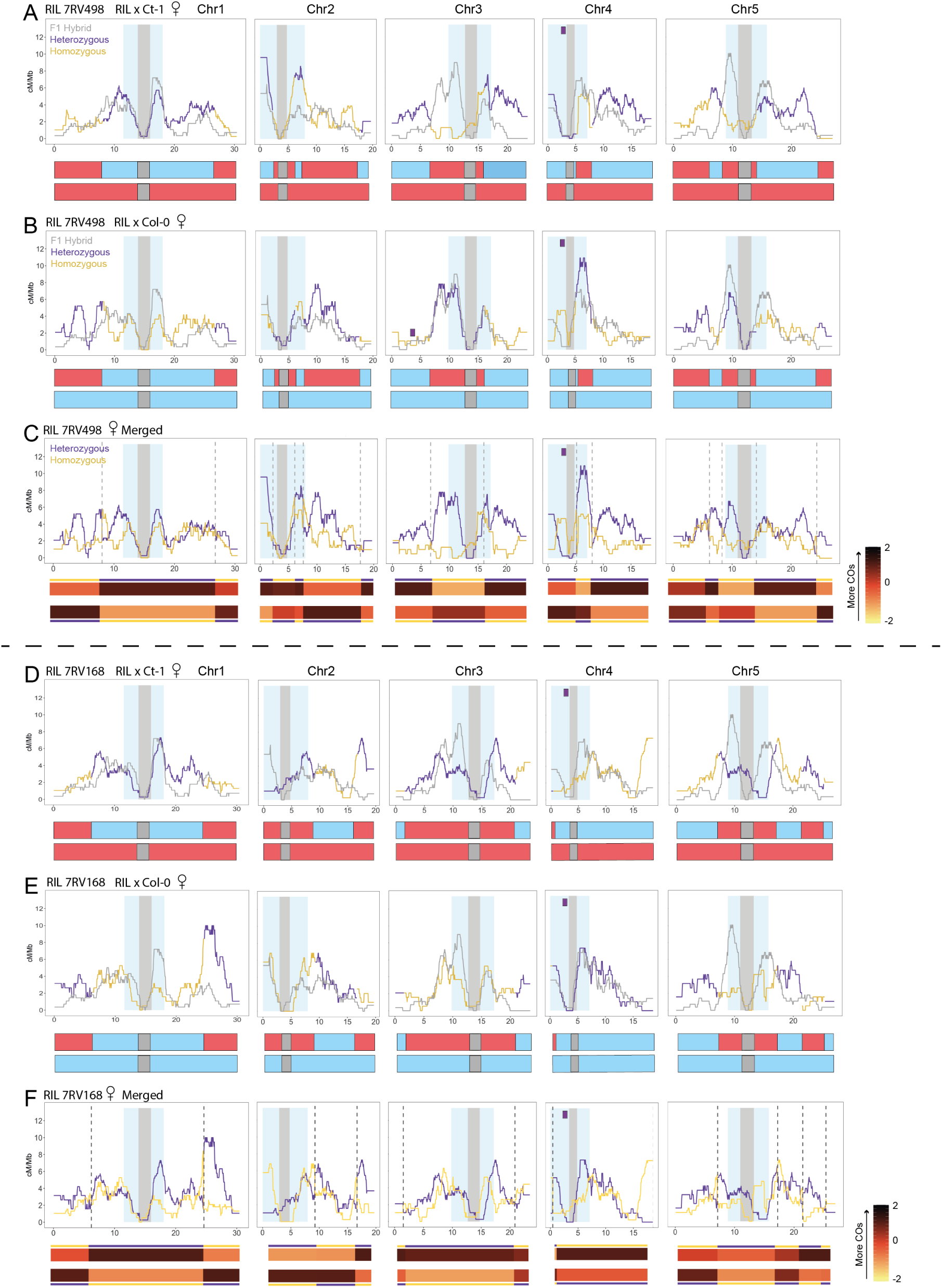
Crossover landscapes in RIL backcrossed female data. **A-C)** Data for female backcross using RIL 7RV498. **A-B)** Each of the reciprocal backcross using the RIL 7RV498 are displayed in the two first panes. **C)** The two reciprocal backcrosses are displayed on the third panel, followed by the heatmap of CO frequency within intervals and each backcross below. **D-F)** Data for female backcross using RIL 7RV168. **D-E)** Each of the reciprocal backcross using the RIL 7RV168 are displayed in the two first panes. **F)** The two reciprocal backcrosses are displayed on the third panel, followed by the heatmap of CO frequency within intervals and each backcross below.

**Fig. S8.**
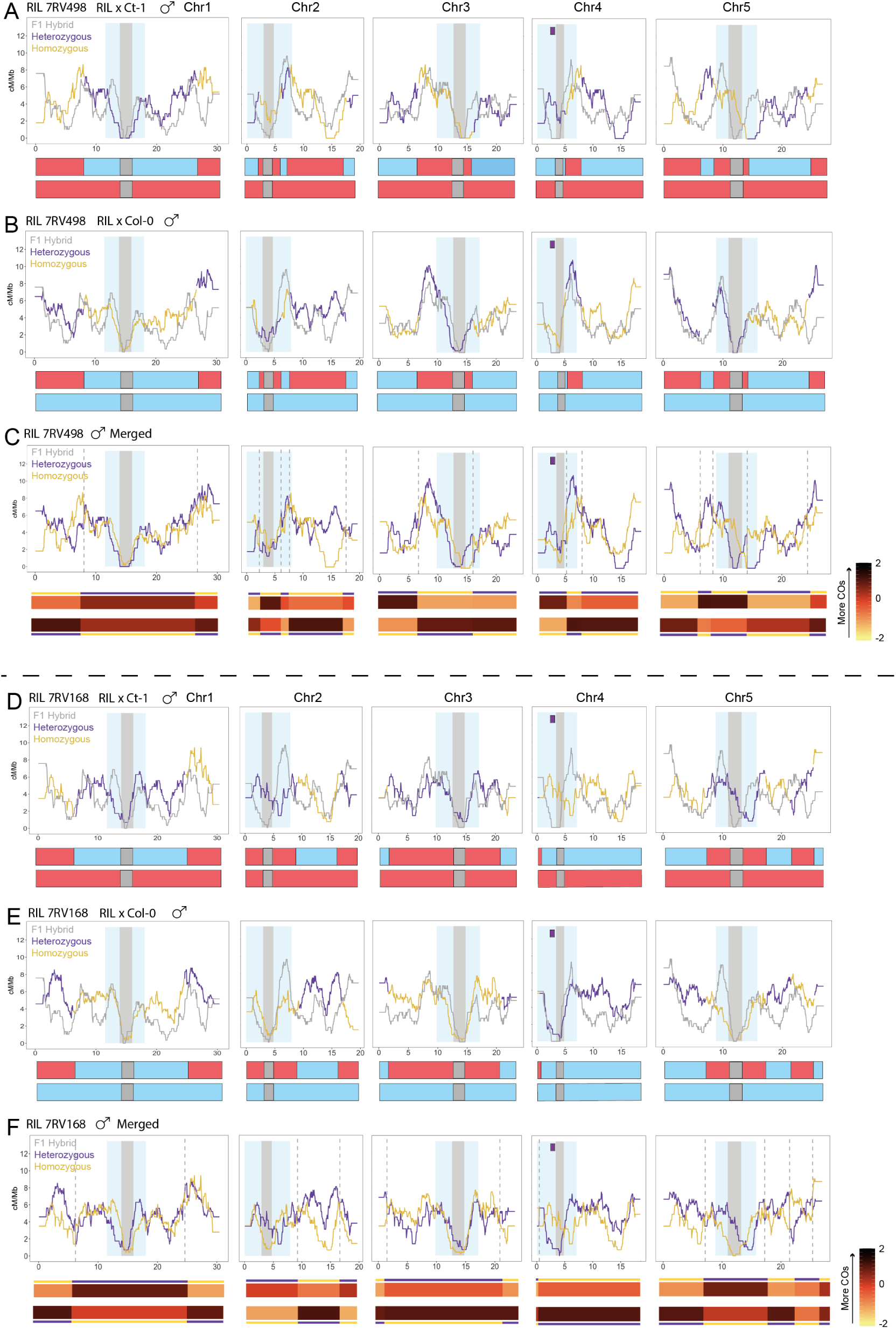
Crossover landscapes in RIL backcrossed male data. **A-C)** Data for female backcross using RIL 7RV498. **A-B)** Each of the reciprocal backcross using the RIL 7RV498 are displayed in the two first panes. **C)** The two reciprocal backcrosses are displayed on the third panel, followed by the heatmap of CO frequency within intervals and each backcross below. **D-F)** Data for male backcross using RIL 7RV168. **D-E)** Each of the reciprocal backcross using the RIL 7RV168 are displayed in the two first panes. **F)** The two reciprocal backcrosses are displayed on the third panel, followed by the heatmap of CO frequency within intervals and each backcross below.

**Fig. S9.**
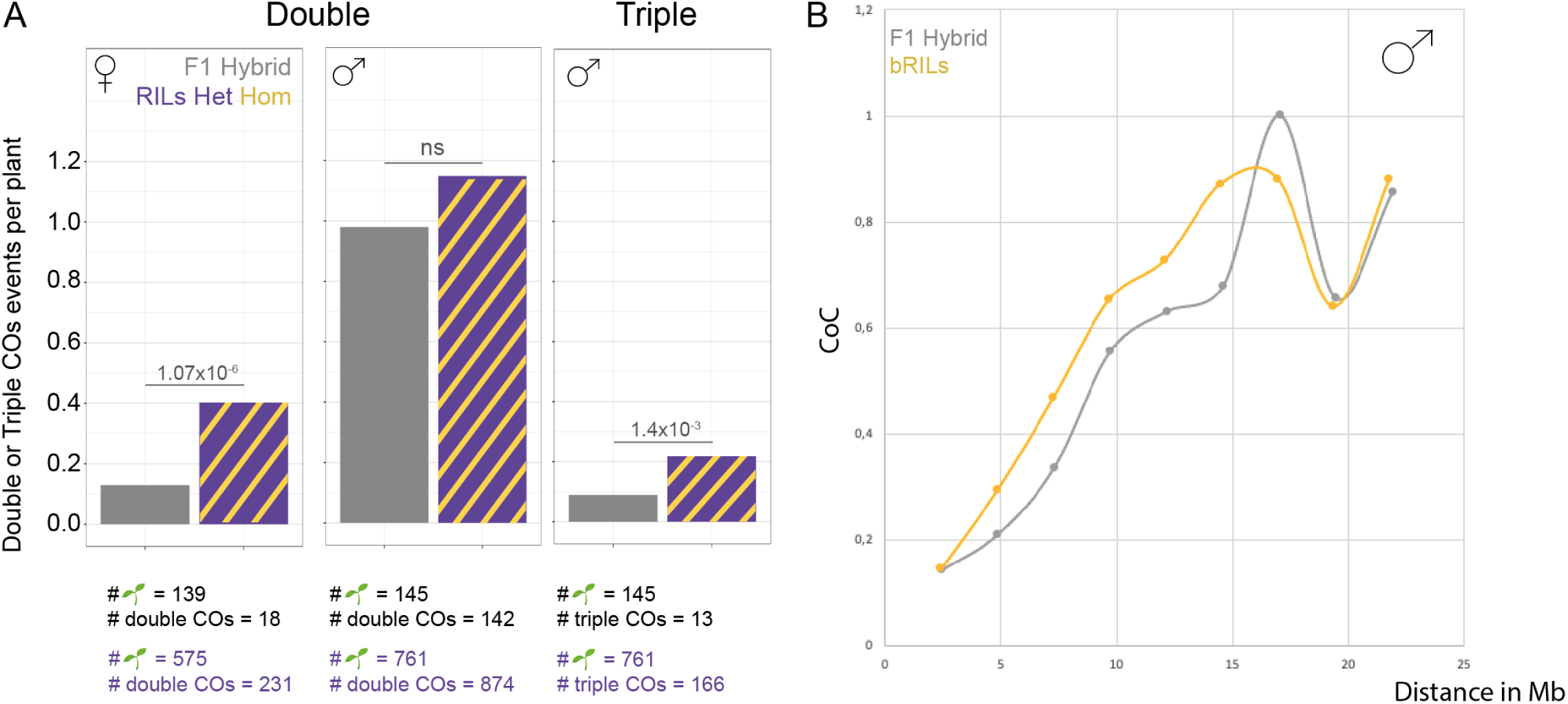
**A)** The number of double crossovers in female and male individuals and triple crossovers in male individuals from F1 hybrid and bRILs populations. Chi-square was used to calculate statistical significance. **B)** Measure of Coefficient of Coincidence in F1 hybrids and bRILs male populations.

**Fig. S10.**
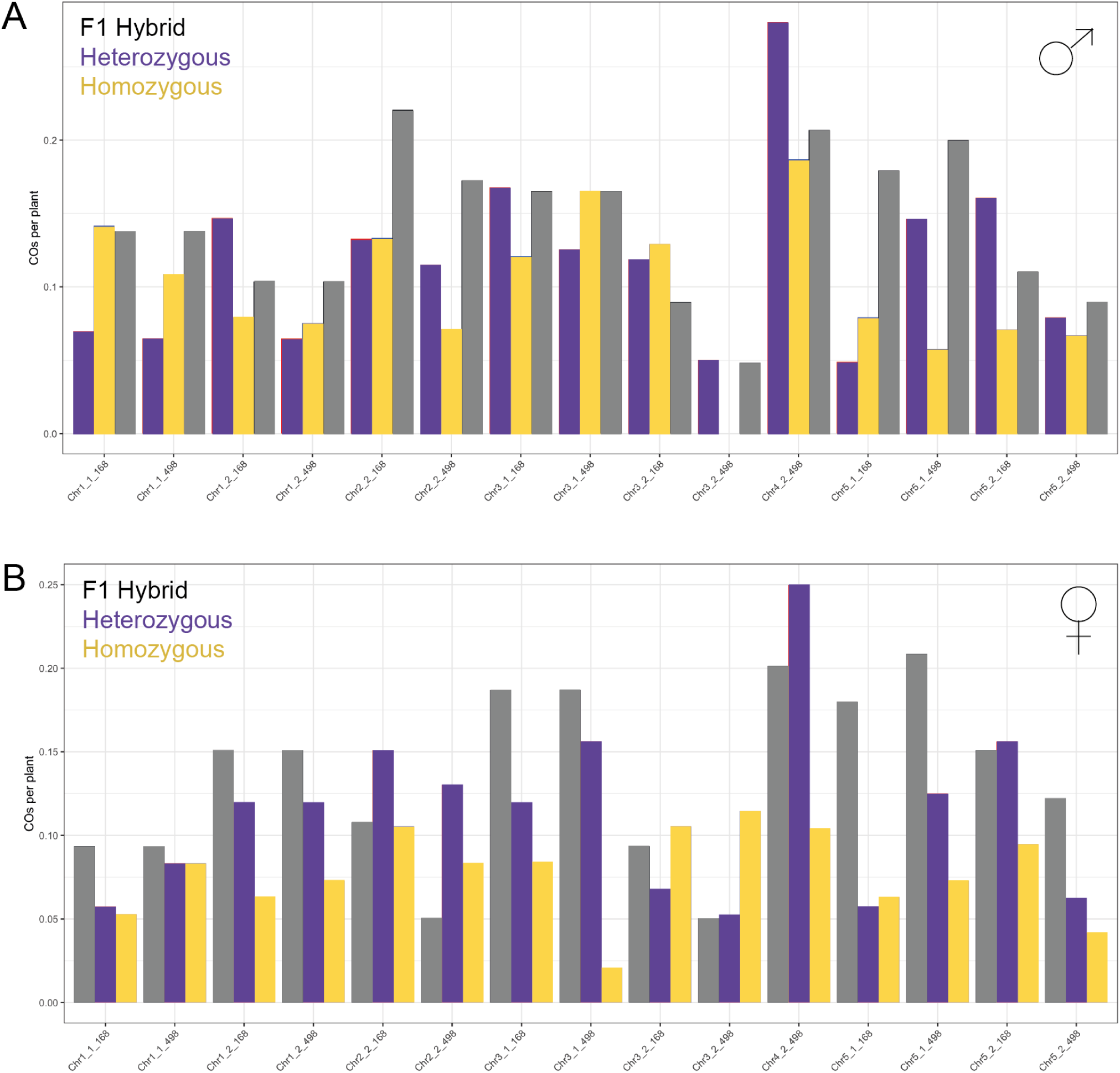
Crossovers per plant within each pericentromeric intervals and interfence measurements. **A-B)** The CO frequency with 15 intervals (S3) was studied for RIL backcrossed individuals and F1 hybrids. Both male and female meiosis were studied.

**Fig. S11.**
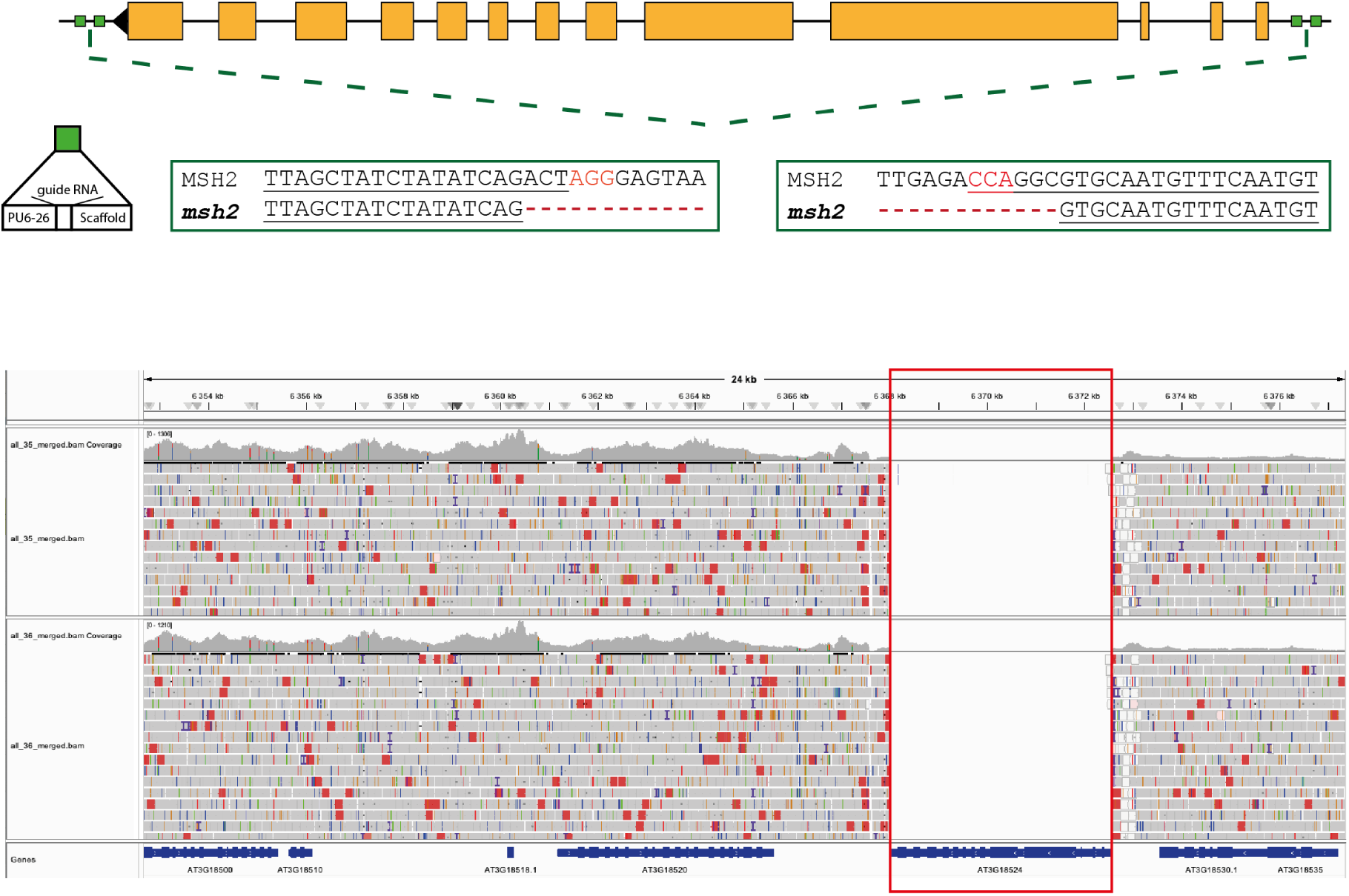
Knock-out MSH2 mutants. Top panel: schematic representation of the MSH2 gene and CRISPR knock-out strategy. Bottom panell: IGV window of 288 individuals sequencing data (two populations from crosses between the RIL 7RV498 with Columbia-0 EMS and Catania-1 EMS). No sequencing read have been mapped to the MSH2 gene, suggesting that no individual of the population possess this gene.

**Fig. S12.**
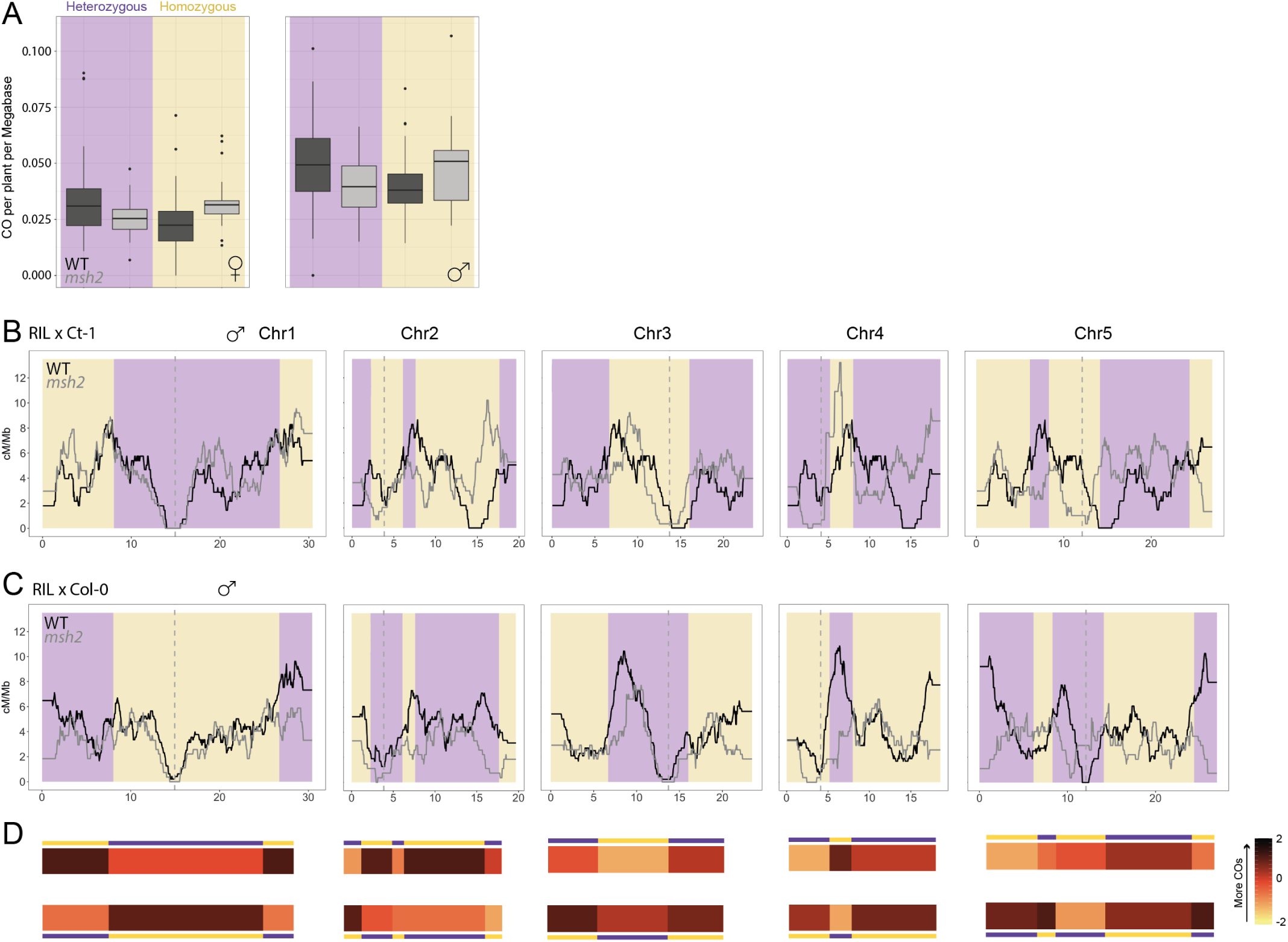
Crossover landscapes in RIL backcrossed male data knock-out mutant *msh2* The two reciprocal backcrosses are displayed followed by the heatmap of CO frequency within intervals below.

**Fig. S13.**
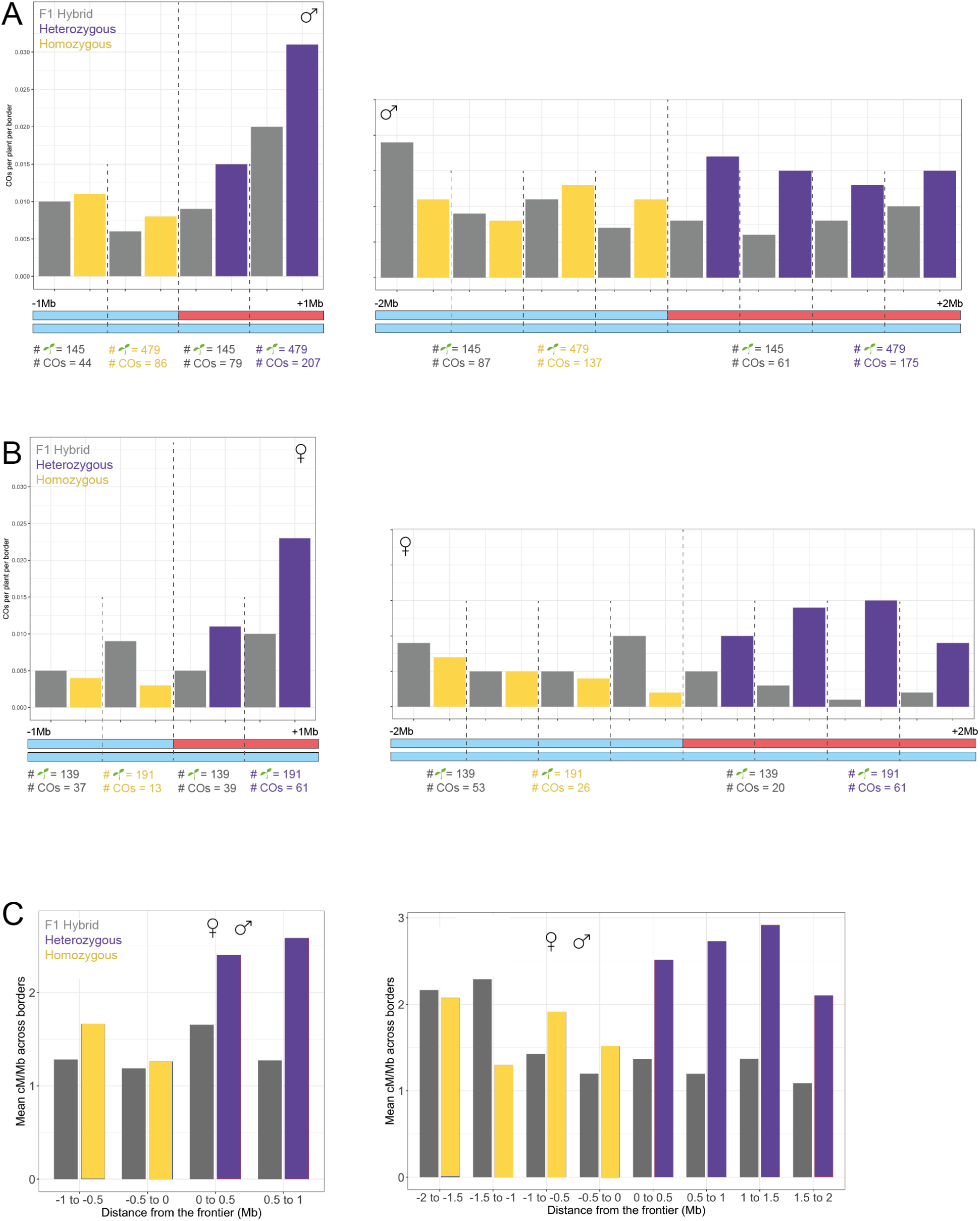
Distribution of crossovers around frontiers. **A-C)** The number of COs across 19 borders within the two RILs, male and female separately. The position of the COs was retrieved 1Mb around the borders and the numbers of events were plotted in four 0.5Mb windows (-1Mb to -0.5Mb, -0.5Mb to the frontier, frontier to 0.5Mb and 0.5Mb to 1Mb). The same analysis was done for 13 intervals of 2Mb around frontiers, yielding eight windows of 0.5Mb. **A-B** This analysis was done using Colombia-0 EMS mutations markers. **C** This analysis was done using Colombia-0 EMS mutations markers for homozygous regions and the Col-0/Ct-1 divergence for heterozygous regions, pooled for both male and female populations.

**Table S1:** Number of EMS markers per population.

| Female x Male | Col-0 EMS | Ct-1 EMS |
| --- | --- | --- |
| I - 498 x Col xx 498 x Ct | 768 | 374 |
| II - 498 x Ct xx 498 x Col | 744 | 433 |
| IV - 168 x Col xx 168 x Ct | 756 | 496 |
| V - 168 x Ct xx 168 x Col | 756 | 509 |
| XV - Col xx 498 x Col | 675 | NA |
| XVI - Col xx 498 x Ct | NA | 339 |
| XVII - Col xx 168 x Col | 669 | NA |
| XVIII - Col xx 168 x Ct | NA | 493 |
| XXVIII - Col x Ct xx Col | 793 | 488 |
| XXX.1 - Col xx Col x Ct | 694 | 447 |
| XXX.2 - Col xx Col x Ct | 672 | 455 |
| XXXV - 498 x Col xx 498 x Ct *msh2* | 673 | 460 |
| XXXVI - 498 x Ct xx 498 x Col *msh2* | 781 | 443 |

**Table S2:**
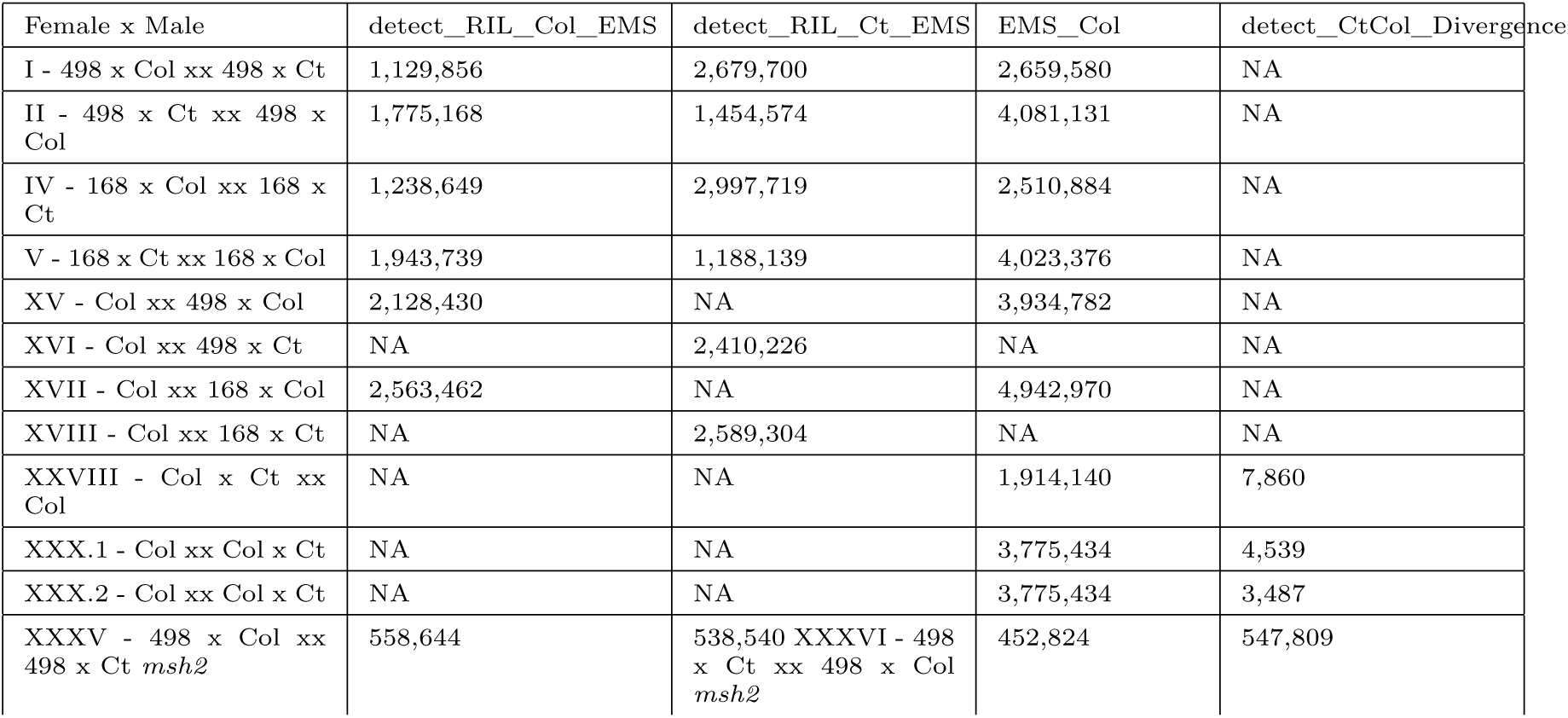
Resolution of crossover detection depending on the sets of markers.

**Table S3:** Number of crossovers detected in F1 hybrid female individuals. Three methods have been used to detect crossovers: the Ct-1/Col-0 divergence, markers of the Col-0 EMS line and markers of the Ct-1 EMS line.

| Method | #COs | #COs/plant |
| --- | --- | --- |
| detect_CtCol_Divergence | 384 | 2.8 |
| EMS_Col | 380 | 2.7 |
| EMS_Ct | 425 | 3.1 |

**Table S4:** Number of crossovers detected in F1 hybrid male individuals. Three methods have been used to detect crossovers: the Ct-1/Col-0 divergence, markers of the Col-0 EMS line and markers of the Ct-1 EMS line.

| Method | #COs | #COs/plant |
| --- | --- | --- |
| detect_CtCol_Divergence | 658 | 4.5 |
| EMS_Col | 621 | 4.3 |
| EMS_Ct | 574 | 4.0 |

**Table S5:** Number of crossovers per individual in F1 hybrids and RIL backcrosses male and female populations.

| Method | Population | COs per individual |
| --- | --- | --- |
| detectCO_ColCt_divergence | F1_Hybrids_Male | 4.57 |
| detect_RIL_EMS | RIL_Backcrosses_Male | 5.32 |
| detectCO_ColCt_divergence | F1_Hybrids_Female | 2.77 |
| detect_RIL_EMS | RIL_Backcrosses_Female | 3.29 |

**Table S6:**
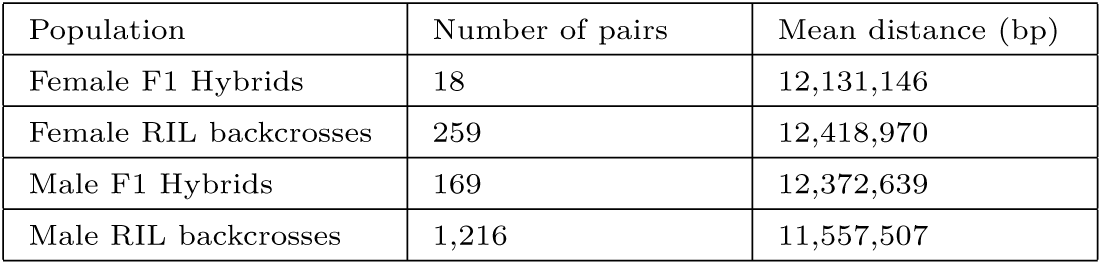
Mean inter_CO distances in all four populations.

| Population | Number of pairs | Mean distance (bp) |
| --- | --- | --- |
| Female F1 Hybrids | 18 | 12,131,146 |
| Female RIL backcrosses | 259 | 12,418,970 |
| Male F1 Hybrids | 169 | 12,372,639 |
| Male RIL backcrosses | 1,216 | 11,557,507 |

**Table S7:**
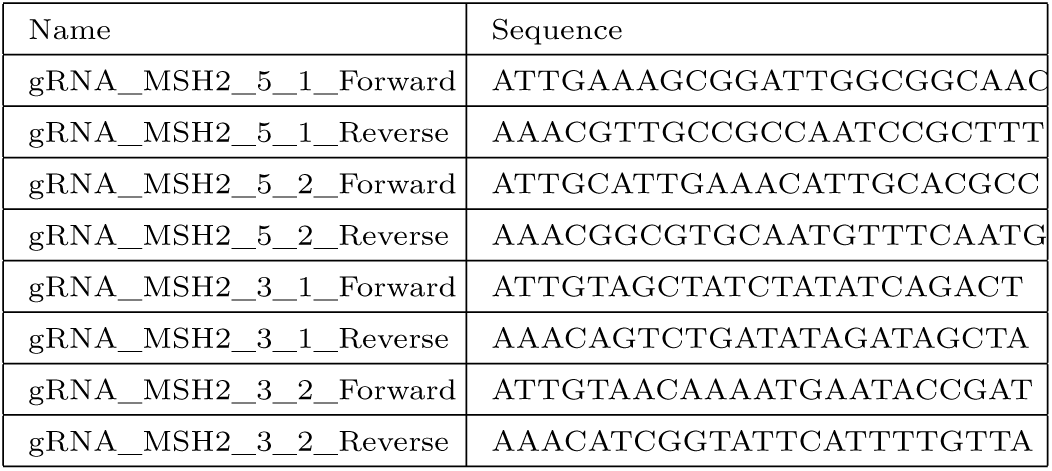
Oligonucleotides used for cloning within CRISPR/Cas9 goldenbraid backbones. A ramp-down from 60°C to 50°C was used to hybridize Forward and Reverse strands together before cloning.

**Table S8:** Plasmids used for engineering *msh2* KO lines in *A. thaliana*.

| Cassette(s) | Backbone | Resistance cassette | Reference |
| --- | --- | --- | --- |
| U6-26 | pUPD | Ampicillin | GB1001 |
| psgRna | pUPD | Ampicillin | GB0645 |
| EMPTY | p_alpha1 | Kanamycin | RpDGB3_alpha1 |
| EMPTY | p_alpha2 | Kanamycin | pDGB3_alpha2 |
| EMPTY | p_omega1 | Spectinomycin | pDGB3_omega1 |
| EMPTY | p_omega2 | Spectinomycin | pDGB3_omega2 |
| Cas9 + DsRed | p_alpha2 | Kanamycin | X |
| Cas9 + DsRed | p_omega2 | Spectinomycin | X |

**Table S9:**
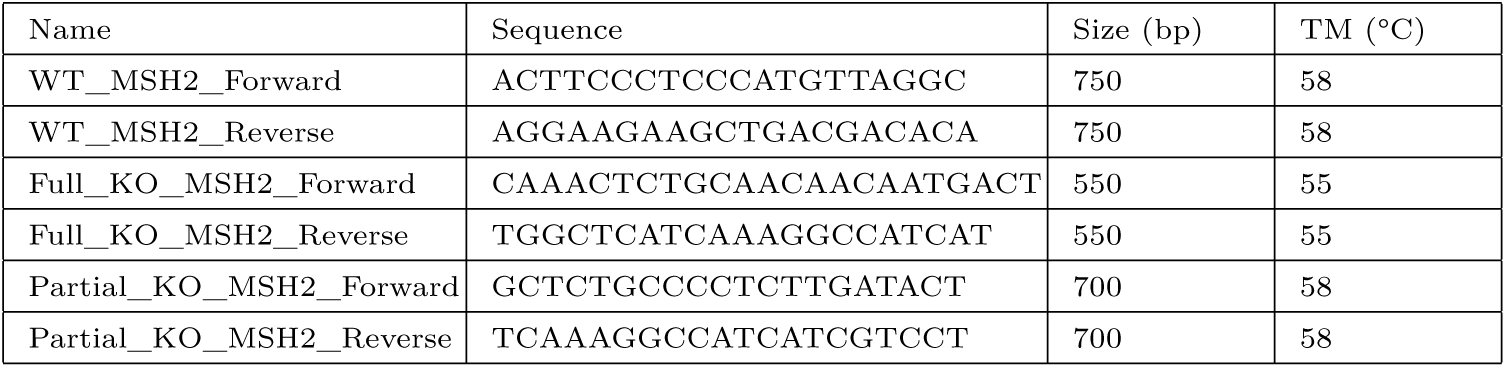
Oligonucleotides used for genotyping the *msh2* deletion.

